# Lysine biosynthesis impairment shapes heat-stress acclimation through metabolic and transcriptional reprogramming in *Arabidopsis thaliana*

**DOI:** 10.64898/2026.09.07.749967

**Authors:** Débora Gonçalves Gouveia, Philipp Westhoff, Wesley E. B. Barrios, Mona Perrar, Samantha Flachbart, Auxiliadora Martins, Pedro A. B. Reis, Adriano Nunes-Nesi, Jürgen Zeier, Andreas P. M. Weber, Wagner L. Araújo

## Abstract

Global warming is increasing the frequency and intensity of high-temperature episodes, limiting plant productivity. However, the molecular mechanisms integrating primary metabolism with the heat stress response remains poorly understood. Here, we show that lysine biosynthesis contributes to the coordination of physiological, metabolic and transcriptional responses to heat stress in *Arabidopsis thaliana*. We compared wild-type, the lysine-biosynthesis mutant *dapat*, and the salicylic acid (SA)-biosynthesis and signaling mutants *sid2-1* and *npr1-3* under prolonged warming (6□°C above control for 7 days) and heat shock (38 °C for 6 h), followed by recovery. We assessed growth, gas exchange, photosynthetic performance, free SA, salicylic acid glucoside (SAG), salicylic acid glucose ester (SGE), and total SA content, primary metabolite profiles, heat-stress-responsive gene expression and transcriptome-wide changes by RNA sequencing. Before heat stress, *dapat* mutant presented a distinct metabolic state, marked by amino-acid accumulation, altered organic-acid profiles, reduced soluble sugars and elevated endogenous SA. This metabolic configuration persisted during prolonged warming, whereas WT and SA-pathway mutants underwent more dynamic reprogramming. Heat shock, by contrast, elicited a more convergent response across genotypes. Despite reduced basal PSII efficiency, *dapat* maintained photosynthetic performance during prolonged warming and recovered. Its transcriptional response, however, differed from that of WT and SA-pathway mutants: selected heat-responsive genes were constitutively or more strongly expressed, whereas some canonical heat- stress regulators showed weaker induction after heat shock. RNA-seq further revealed a largely conserved core heat-shock response but genotype-dependent regulation of defense, hormone and amino-acid-metabolism programs, particularly during recovery. Together, these findings indicate that impaired DAPAT activity establishes a metabolically primed but energetically constrained state that reshapes gas exchange, photosynthetic acclimation and heat-responsive transcription. Lysine homeostasis therefore emerges as a regulatory node linking primary metabolism and SA accumulation with SA-dependent and SA-independent components of heat-stress acclimation.

## 1. Introduction

Anthropogenic climate change is causing unprecedented thermal stress on terrestrial ecosystems, with global mean temperatures projected to increase by at least 1.5°C by 2040 and extreme heat events becoming increasingly frequent and severe (Shukla *et al*., 2022). These thermal perturbations impose critical constraints on plant productivity and agricultural sustainability, as temperatures exceeding optimal growth ranges by 10-15°C trigger cascading cellular dysfunction including excessive reactive oxygen species (ROS) accumulation, protein denaturation, membrane destabilization, and irreversible DNA damage (Jagadish *et al*., 2021). Such thermal stress fundamentally disrupts photosynthetic machinery, carbon assimilation pathways, hormonal homeostasis, and reproductive development, ultimately culminating in cellular death and substantial yield reductions. To mitigate these deleterious effects, plants have evolved sophisticated adaptive mechanisms encompassing heat shock protein (HSP) synthesis, heat-responsive (HSR) genes activation, antioxidant defense system mobilization, and comprehensive metabolic reprogramming that collectively enhance thermotolerance (Jacob *et al*., 2017; Sato *et al*., 2024). These adaptive responses are coordinated by intricate transcriptional networks that define specific branches of the heat stress response pathway and determine response specificity under elevated temperatures (Ding *et al*., 2020; Haider *et al*., 2022; Liu *et al*., 2025).

Central to this regulatory module is salicylic acid (SA), a phytohormone that functions as a key signaling hub integrating developmental processes with responses to biotic and abiotic challenges (Khan *et al*., 2013; Fayez & Bazaid, 2014; Khan *et al*., 2014; Nazar *et al*., 2015; Zhang *et al*., 2015). While SA is predominantly recognized for its role in systemic acquired resistance through pathogen defense priming (Klessing *et al*., 2018), emerging evidence demonstrates its pivotal function in thermotolerance through HSP accumulation enhancement, ROS scavenging facilitation, and photosynthetic machinery preservation (Khan *et al*., 2013; Jahan *et al*., 2019; Sangwan *et al*., 2022). SA biosynthesis in plants occurs primarily through the isochorismate synthase pathway, with ISOCHORISMATE SYNTHASE2 (ICS2/SID2; EC 4.4.2) serving as the key enzyme catalyzing SA production (Wildermuth *et al*., 2001; Garcion *et al*., 2008), while SA perception and signaling are mediated through the NPR1 protein containing ankyrin repeats domain that controls systemic acquired resistance responses (Cao *et al*., 1997; Wu *et al*., 2012). Both exogenous SA application and elevated endogenous SA levels significantly enhance basal thermotolerance, with *Arabidopsis*-SA mutants exhibiting compromised heat shock survival while SA-accumulating genotypes demonstrate superior resilience during thermal stress and subsequent recovery phases, though notably, SA signaling appears dispensable for acquired thermotolerance following heat acclimation (Clarke *et al*., 2004). The metabolic regulation of SA homeostasis involves complex conjugation mechanisms, including glucose conjugations mediated by UGT74F1 and UGT74F2 enzymes that modulate SA availability and signaling intensity (Thompson *et al*., 2017).

Lysine has emerged as a focal point in plant metabolic research over the past four decades, with its critical role in stress responses becoming increasingly apparent through comprehensive biochemical and molecular investigations (Hildebrandt, 2018; Yang *et al*., 2020; Singh *et al*., 2026). While lysine biosynthesis pathways are conserved across prokaryotic and eukaryotic organisms, significant mechanistic divergences exist between taxonomic groups, with plant systems utilizing the diaminopimelate (DAP) pathway wherein the pyridoxal phosphate-dependent enzyme *L,L*-diaminopimelate aminotransferase (DAPAT; EC 2.6.1.83) catalyzes a critical transamination reaction (Hudson *et al*., 2005, 2006).

Beyond its biosynthetic role, lysine undergoes extensive degradation processes that generate alternative metabolic substrates crucial for stress adaptation, with lysine catabolism producing glutamate, 2-oxoglutarate (2-OG) and acetyl-CoA that feed into central metabolic pathways including the TCA cycle and the mETC (Hildebrandt *et al*., 2015; Arruda & Barreto, 2020). Under heat stress conditions, plants exhibit enhanced reliance on alternative respiratory pathways and amino acid catabolism, with branched-chain amino acids (BCAA) serving as primary substrates for energy production and stress tolerance mechanisms (Bulut *et al*., 2025). The metabolic interconnection between lysine and BCAA metabolism becomes particularly relevant under heat stress, as both amino acid pools serve as alternative substrates for maintaining cellular energy homeostasis when conventional photosynthetic and respiratory processes are compromised (Araújo *et al*., 2010, 2011; Hildebrandt, 2018; Heinemann & Hildebrandt, 2021).

The *Arabidopsis dapat* mutant carries a point mutation that reduces *L,L*-diaminopimelate aminotransferase (*DAPAT*) activity to approximately 10% of wild-type levels. This defect causes growth inhibition, altered leaf architecture and extensive metabolic reprogramming, and modifies responses to environmental stress (Cavalcanti *et al*., 2018; Neves *et al*., 2024; Gouveia *et al*., 2025, *bioRxiv preprint*; Gouveia *et al*., 2026). Under control conditions, *dapat* plants also accumulate free salicylic acid (SA) and display enhanced SA-dependent resistance to *Pseudomonas syringae*, associated with increased expression of pathogenesis-related genes (Rate & Greenberg, 2001; Song *et al*., 2004).

These phenotypes are consistent with a chronic stress-like metabolic state characterized by energy limitation, altered carbon–nitrogen balance, amino-acid accumulation, changes in branched-chain amino-acid metabolism and reduced sugar availability (Cavalcanti et al., 2018; Gouveia et al., 2026). This state is supported by increased reliance on alternative mitochondrial respiratory pathways and amino-acid catabolism (Araújo *et al*., 2010, 2011; Cavalcanti *et al*., 2018; Gouveia *et al*., 2025, *bioRxiv preprint*). Together with constitutively elevated SA, the metabolic consequences of *DAPAT* deficiency suggest that lysine biosynthesis may influence heat-stress responses through both metabolic and signaling mechanisms. Whether this pre-existing state enhances or constrains thermotolerance, however, remains unknown.

To address this question, we compared wild-type (WT), the lysine-biosynthesis- deficient *dapat* mutant, and the SA-biosynthesis and signaling mutants *sid2-1* and *npr1-3* under prolonged warming and heat shock followed by recovery. We integrated physiological measurements, SA quantification, primary-metabolite profiling and RNA- seq to assess how lysine biosynthesis affects gas exchange, photosynthetic performance, heat-responsive transcription and metabolic recovery. We found that *DAPAT* deficiency does not simply enhance or compromise heat tolerance. Instead, it establishes a metabolically primed but energetically constrained state, characterized by elevated basal SA, persistent amino-acid accumulation, altered carbon metabolism and stomatal regulation, dynamic reconfiguration of SA pools, and distinct transcriptional regulation of heat-, defense- and recovery-associated programmes. Although *dapat* plants maintained core photosynthetic performance and activated the canonical heat-shock response, their broader metabolic and transcriptional responses differed from those of WT and SA-pathway mutants, particularly during recovery. These findings position lysine biosynthesis as a regulatory node that links primary metabolism with SA homeostasis and heat-stress transcription, thereby shaping the balance between physiological resilience and metabolic cost during heat acclimation.

## 2. Materials and methods

### 2.1. Plant material

*Arabidopsis* wild type (WT), *dapat* - a lysine biosynthesis mutant (Rate & Greenberg, 2001; Cavalcanti *et al*., 2018; Gouveia *et al*., 2026), *sid2-1* (Garcion et al., 2008; Zhang et al., 2010) and *nrp1-3* (Cao et al., 1997) mutant lines, for salicylic acid biosynthesis and signaling, respectively were used. The seeds were surface sterilized by immersion in 1 mL of 80% (v/v) ethanol containing 0.05% (v/v) Tween 20 for 5 minutes under constant stirring. Subsequently, the ethanol solution was removed, and the seeds were subjected to an additional 5-minute wash with 100% ethanol under constant stirring. Residual ethanol was removed, and the seeds were air-dried for 2 hours under sterile conditions. After sterilization, the seeds were aseptically transferred to Petri plates containing half-strength Murashige and Skoog (MS) basal medium (Murashige & Skoog, 1962), supplemented with 1% (w/v) sucrose. The plates were incubated in darkness at 4°C for 72 hours to promote stratification. After two weeks on plates, seeds germinated, and seedlings were moved into pots.

### 2.2. Growth under short-day conditions

*Arabidopsis* wild-type and *dapat* mutant plants were grown under short-day conditions (SD, 8h light/16h dark) at 20°C (day/night), 150 μmol photons m ²s ¹ light intensity, and 60% of relative humidity.

### 2.3. Prolonged warming and Recovery period

For the prolonged warming treatment, the temperature increased by 6 °C above control (28°C day/night). 21-day-old plants (on pots) were exposed to treatment for a period of 7 days. The whole rosette was harvested in the middle of day, on day 7, for subsequent metabolic and molecular analyses. 28-day-old *Arabidopsis* plants grown under a 8-hour photoperiod followed by prolonged warming, were subjected to a 7-day recovery phase under initial control conditions (20°C day/night). The whole rosette was harvested in the middle of day.

#### 2.3.1. Heat shock treatment

Since 45 °C is considered a lethal temperature for plants, and 37 - 42 °C is a range commonly used in heat stress studies, the sublethal temperature of 38 °C for 6 hours was stipulated as the heat shock (HS) treatment (Wang *et al*., 2020; Réthoreté *et al*., 2024). In the control treatment, the first set of plants were grown under short-day conditions (SD, 8h light/16h dark) at 20°C (day/night). For the heat shock treatment, the temperature was kept at 38°C, with 28-d ay-old plants (on pots) exposed to treatment for 6h. The whole rosette was harvested in the middle of day.

#### 2.3.2. Growth under neutral-day conditions

Plants Col-0, *dapat*, *sid2-1* and *npr1-3* mutants were grown under neutral-day conditions (12h light/12h dark) with temperature variation (22°C day/20°C night), 150 μmol photons m□² s□¹ light intensity, and 60% of relative humidity.

### 2.4. Prolonged warming treatment

Plants were allocated under a 12-hour photoperiod (ND) with day/night temperatures maintained at 22/20□°C (control conditions). For the prolonged warming treatment, the temperature increased by 6 °C above control (28°C day/26°C night). 14- day-old plants (on pots) were exposed to the treatment for a period of 7 days. The whole rosette was harvested in the middle of day, on day 0, 2, 4 and 7 after treatment, for subsequent metabolic and molecular analyses.

### 2.5. Heat shock and Recovery period

For the heat shock treatment, the temperature was kept by 38°C, with 21-day-old plants (on pots) exposed to treatment for 6h. The whole rosette was harvested in the middle of day. 21-day-old *Arabidopsis* plants grown under a 12-hour photoperiod followed by heat shock, were subjected to a 24-h recovery phase. Tissue samples were collected at midday, 6 hours (immediately after heat shock), and 30 hours (24 hours into recovery) for biochemical and molecular analyses.

### 2.6. Measurements of photosynthetic parameters

Measurements were performed with an open-flow infrared gas-exchange analyzer system (Li-Cor 6400XT) with a portable photosynthesis system, in full rosettes of 4- week-old plants maintained in PW and upon HS recovery. *Fv/Fm*, which corresponds to the potential quantum yield of the photochemical reactions of PSII and represents a measure of photochemical efficiency, was measured as described previously (Oh *et al*., 1996) during dark treatment.

### 2.7. Metabolite Profiling

For metabolite profiling, 1.5 mL of pre-cooled (−20°C) mixture of H2O:MeOH:CHCl3 (1:2.5:1 v:v) was used, containing 50 μL / 50 mL of the 5 mM Ribitol/DMPA stock solution (Ribitol and DMPA as internal standards) into each sample and mixed by vortex for 20 sec. Samples were shaken on an orbital shaker for 10 min at 4°C. The samples were then centrifuged at 20,000 g for 5 min at 4°C. After centrifugation, 1.0 mL of supernatant was carefully aspirated and transferred to a clean 1.5-ml microcentrifuge tube. Samples were stored at -80 °C until derivatization for GC-MS.

For GC-MS analysis 30 μl of sample was freeze dried using speed-vac. Dried samples were derivatized (Gu *et al*., 2012; Shim *et al*., 2019). Raw data were converted to the mzXML format using ProteoWizard and to the NetCDF format via MetAlign using default parameters. Deconvolution of mass spectra was conducted using the free deconvolution software AMDIS (Automated Mass Spectral Deconvolution and Identification System from NIST). Deconvoluted mass spectra were matched against the NIST14 Mass Spectral Library (https://www.nist.gov/srd/niststandard-reference-database-1a-v14). Database matches with more than 70% were further compared with an in-house chemical standard library for compound annotation. Compounds, that could not be verified by the in-house library, are named according to the matched compound class and the retention time. Extracted peaks were integrated using MassHunter Quantitative (v b08.00, Agilent Technologies). All metabolite peak areas were normalized to tissue fresh weight and adjusted by the Ribitol internal standard peak area (Sigma-Aldrich) to account for technical variability.

### 2.8. Determination of SA and related compounds

At the indicated times after prolonged warming and heat shock treatments, three leaves from one plant were harvested, combined for one biological replicate, and shock frozen in liquid nitrogen. The levels of SA, SAG, and SA glucose ester (SGE) were determined as detailed in Yildiz *et al*. (2021) and Hartmann *et al*. (2018). In summary, ∼50 mg of pulverized, frozen leaf sample was extracted twice with 1mL of MeOH/50mM sodium phosphate (pH 6.0) (80:20, v/v). In the first extraction step, the extraction buffer was supplemented with a mix of different internal standards (1 μg each). Aliquots of 600 μL of the combined extracts were evaporated to dryness in a Scan Speed vacuum centrifuge (Labogene ApS), and analytes derivatized with 20 μL N□methyl□N□trimethyl□ silyltrifluoroacetamide (MSTFA) containing 1% trimethylchlorosilane (v/v) in 20 μL pyridine and 60 μL of hexane for 30 min at 70°C. Two microliters of the sample mixture was analyzed by gas chromatography mass spectrometry (GCoroacetamide (MSTFA)MS) with an Agilent 7890A/5975C GC□MS system equipped with a Phenomenex ZB□35 (30 m × 0.25 mm × 0.25 μm) capillary column and MSD ChemStation software version E.02.01.1177 (Agilent Technologies). The temperature of the GC oven was programmed as follows: 70°C for 2 min, gradient to 250°C at 10°C/min, gradient to 260°C at 2°C/min, gradient to 320°C at 10°C/min, and 320°C for 3 min. For the quantitative determination of metabolites, specific selected ion chromatograms of analytes and related internal standards were integrated [analyte (m/z) related to internal standard (m/z)]: SA (m/z 267) related to D4□SA (m/z 271); SAG (m/z 267) and SGE (m/z 193) were all related to salicin (m/z 268). For absolute quantification, experimental correction actors determined via GC□MS analysis of authentic compounds were considered, and metabolite levels were related to the fresh weight of the leaf samples (Yildiz *et al*., 2023).

### 2.9. Gene Expression Analysis

Total RNA was extracted from plant tissues using the RNeasy Plant Mini Kit (Qiagen, Cat. No. 74904) with on-column DNase I treatment (RNase-free, New England Biolabs, Cat. No. M0303) to eliminate genomic DNA contamination. RNA concentration was determined using a NanoDrop OneC Microvolume UV-Vis Spectrophotometer (Thermo Fisher Scientific), with A260/A280 ratios >2.0 and A260/A230 ratios >1.8 indicating high-quality RNA. RNA integrity was verified by 1.5% agarose gel electrophoresis stained with GelRed (Biotium), visualizing distinct 28S and 18S ribosomal RNA bands. First-strand cDNA synthesis was performed using the LunaScript RT SuperMix Kit (New England Biolabs, Cat. No. E3010) in 20 μL reactions. The reverse transcription protocol consisted of 25°C for 2 min, 55°C for 10 min, and 95°C for 1 min. No-RT controls were included to confirm absence of genomic DNA contamination.

Quantitative PCR was performed using Luna Universal qPCR Master Mix (New England Biolabs, Cat. No. M3003) on an Applied Biosystems Step OnePlus Real-Time PCR System. Thermal cycling conditions were 95°C for 60 s, followed by 40 cycles of 95°C for 15 s and 60°C for 30 s, with melting curve analysis from 60°C to 95°C.

Gene expression was analyzed using the comparative CT method (ΔΔCT) with *UBC9*, *UBC21* and *PP2A* as reference genes. All samples were analyzed in biological triplicates, and primer efficiencies were validated using serial dilutions (90-110% efficiency, R² >0.99). Primers were designed using Primer3Plus web application. Statistical analysis was performed using R Studio software with pre-calculated 2^(-ΔΔCT)^ values and appropriate statistical testing.

### 2.10. Experimental design and statistical analysis

Statistical analyses for Table 1; Figures 2a,b,d, 4–7; and Supplementary Figs. S2a,b,d and S4-5 were performed using two-way ANOVA followed by Tukey’s HSD test for multiple comparisons (α = 0.05). Different letters indicate statistically significant differences among groups. For Figure 2c and Supplementary Fig. S2c, pairwise comparisons with the WT control were performed using Student’s t-test. Statistical significance is indicated by an asterisk conventions (*p < 0.05, ***p<0.001) or value, whereas non-significant comparisons are left unmarked.

**Table 1.** Gas-exchange and photosynthetic parameters of *Arabidopsis* plants during prolonged warming, heat shock and recovery. The *dapat* mutant was used to investigate defects in lysine biosynthesis, whereas the *sid2-1* and *npr1-3* lines were used to examine disruptions in salicylic acid biosynthesis and signaling, respectively. Gas- exchange and photosynthetic parameters were measured in 5-week-old wild-type, *dapat, sid2-1* and *npr1-3* plants grown under neutral-day conditions at control temperatures (22 °C during the day and 20 °C at night). For prolonged warming, day and night temperatures were increased by 6 °C for 7 days. For heat shock, plants were exposed to the treatment and subsequently evaluated after 24-h of recovery. The measured parameters included stomatal conductance (*gs*), apparent transpiration rate (*E*), leaf vapor-pressure deficit (VPD*leaf*), effective quantum yield of photosystem II (Φ*PSII*) and electron transport rate (*ETR*). Data are presented as means ± s.e. (n = 8); three leaves per plant were used to calculate each biological replicate. Different letters indicate significant differences according to ANOVA followed by Tukey’s multiple- comparisons test (P < 0.05).

| Genotype | Treatment | Stomatal conductance (mol m<sup>-2</sup> s<sup>-1</sup>) | Transpiration (mmol m<sup>-2</sup> s<sup>-1</sup>) | Leaf VPD (kPa) | ΦPSII | Electron Transport Rate (μmol m<sup>-2</sup> s<sup>-1</sup>) |
| --- | --- | --- | --- | --- | --- | --- |
| WT | Control | 0.215 ± 0.003<sup>ab</sup> | 1.656 ± 0.049<sup>de</sup> | 0.811 ± 0.024<sup>d</sup> | 0.763 ± 0.003<sup>a</sup> | 2.713 ± 0.146<sup>b</sup> |
| *dapat* | Control | 0.182 ± 0.010<sup>b</sup> | 1.803 ± 0.074<sup>ab</sup> | 1.043 ± 0.019<sup>b</sup> | 0.757 ± 0.002<sup>ab</sup> | 2.087 ± 0.128<sup>c</sup> |
| *npr1-3* | Control | 0.223 ± 0.014<sup>a</sup> | 2.279 ± 0.144<sup>a</sup> | 1.064 ± 0.011<sup>a</sup> | 0.768 ± 0.002<sup>a</sup> | 1.921 ± 0.108<sup>c</sup> |
| *sid2-1* | Control | 0.213 ± 0.010<sup>ab</sup> | 1.999 ± 0.063<sup>ac</sup> | 0.993 ± 0.012<sup>c</sup> | 0.759 ± 0.003<sup>ab</sup> | 2.057 ± 0.119<sup>c</sup> |
| WT | Warming | 0.125 ± 0.007<sup>c</sup> | 1.105 ± 0.046<sup>d</sup> | 0.909 ± 0.014<sup>c</sup> | 0.757 ± 0.003<sup>ab</sup> | 3.642 ± 0.119<sup>a</sup> |
| *dapat* | Warming | 0.132 ± 0.008<sup>c</sup> | 1.301 ± 0.077<sup>de</sup> | 1.070 ± 0.013<sup>a</sup> | 0.763 ± 0.004<sup>a</sup> | 2.138 ± 0.140<sup>c</sup> |
| *npr1-3* | Warming | 0.132 ± 0.004<sup>c</sup> | 1.377 ± 0.037<sup>de</sup> | 1.071 ± 0.007<sup>a</sup> | 0.760 ± 0.001<sup>ab</sup> | 3.369 ± 0.119<sup>a</sup> |
| *sid2-1* | Warming | 0.128 ± 0.007<sup>c</sup> | 1.322 ± 0.071<sup>d</sup> | 1.060 ± 0.011<sup>a</sup> | 0.750 ± 0.003<sup>b</sup> | 3.678 ± 0.120<sup>a</sup> |
| WT | Control | 0.284 ± 0.005<sup>a</sup> | 2.514 ± 0.071<sup>de</sup> | 1.011 ± 0.029<sup>e</sup> | 0.771 ± 0.003<sup>de</sup> | 2.690 ± 0.104<sup>c</sup> |
| *dapat* | Control | 0.220 ± 0.010<sup>b</sup> | 3.070 ± 0.124<sup>a</sup> | 1.485 ± 0.018<sup>a</sup> | 0.758 ± 0.003<sup>abc</sup> | 2.177 ± 0.086<sup>d</sup> |
| *npr1-3* | Control | 0.301 ± 0.017<sup>a</sup> | 3.505 ± 0.127<sup>b</sup> | 1.302 ± 0.023<sup>c</sup> | 0.766 ± 0.003<sup>ade</sup> | 3.250 ± 0.055<sup>b</sup> |
| *sid2-1* | Control | 0.268 ± 0.010<sup>a</sup> | 3.117 ± 0.076<sup>ab</sup> | 1.279 ± 0.009<sup>b</sup> | 0.772 ± 0.001<sup>d</sup> | 3.169 ± 0.121<sup>bc</sup> |
| WT | Recovery | 0.192 ± 0.009<sup>bc</sup> | 2.012 ± 0.071<sup>c</sup> | 1.146 ± 0.005<sup>c</sup> | 0.746 ± 0.001<sup>b</sup> | 2.250 ± 0.161<sup>a</sup> |
| *dapat* | Recovery | 0.178 ± 0.005<sup>c</sup> | 2.115 ± 0.068<sup>c</sup> | 1.256 ± 0.013<sup>b</sup> | 0.750 ± 0.004<sup>bc</sup> | 2.266 ± 0.084<sup>a</sup> |
| *npr1-3* | Recovery | 0.208 ± 0.006<sup>bc</sup> | 2.742 ± 0.068<sup>ad</sup> | 1.395 ± 0.019<sup>d</sup> | 0.761 ± 0.002<sup>ade</sup> | 4.213 ± 0.129<sup>b</sup> |
| *sid2-1* | Recovery | 0.195 ± 0.006<sup>bc</sup> | 2.342 ± 0.065<sup>ce</sup> | 1.330 ± 0.009<sup>bd</sup> | 0.760 ± 0.002<sup>ace</sup> | 3.365 ± 0.132<sup>b</sup> |

### 2.11. RNAseq of Heat-Shock and recovery experiment

Raw single-end RNA-seq reads were subjected to quality control and adapter trimming using fastp, retaining reads with a minimum Phred quality score of 20 and a minimum length of 30 bp. Cleaned reads were aligned to the *Arabidopsis thaliana* TAIR10 reference genome using STAR, with gene-level read counts generated from the corresponding GTF annotation. Alignments were retained as coordinate-sorted BAM files and indexed using SAMtools. Sequencing and alignment quality metrics were summarized using MultiQC. STAR-generated unstranded gene counts were used to construct the gene-by-yielding least-squares mean estimates adjusted for the other model factors. Subsequent pairwise genotype comparisons within each Day×Treatment stratum was performed using Tukey’s honestly significant difference (HSD) test through the cld() function in emmeans, thus controlling the family-wise error rate at α = 0.05 for each set of contrasts. Finally, the pattern of significant differences was summarized as a compact letter display (CLD) using the multcompView package, wherein genotypes sharing the same letter within a given Day×Treatment condition does not differ significantly, and distinct letters denote statistically meaningful differences at the 5% level.

RNA-seq count data were analyzed in R 4.5.0 using DESeq2 1.48.2. The dataset comprised WT and *dapat* plants sampled under Control Heat, Heat Shock, Control Recovery and Heat Recovery conditions, with 3 biological replicates. Genes with fewer than 10 counts in at least three samples were excluded, and library-size normalization was performed using the DESeq2 median-of-ratios method.

For exploratory analyses, variance-stabilizing transformation (VST; blind = FALSE) was applied. Principal component analysis (PCA) was performed using the 1,000 most variable genes, and sample-to-sample distances were used for hierarchical clustering to assess overall transcriptomic relationships and replicate consistency. Hierarchical clustering of the 100 most variable genes was additionally performed after row-wise Z- score normalization.

Differential expression was assessed using a model in which each genotype– treatment combination was represented as a separate group. Log_2_ fold changes were shrunk using ashr where applicable. Genes with FDR < 0.05 and |log_2_ FC| ≥ 0.585 (|FC| ≥ 1.5) were considered differentially expressed. Heat-shock and recovery responses were evaluated separately within each genotype, and genotype differences were assessed within each condition. Genotype-dependent differences in response magnitude were quantified as ΔLFC = LFC(*dapat*) − LFC(WT), with genes additionally classified according to the direction and magnitude of their responses.

Functional enrichment was assessed using Gene Ontology (GO) Biological Process and KEGG over-representation analyses implemented in clusterProfiler. GO terms were tested using a hypergeometric test and Benjamini–Hochberg correction, and redundant terms were reduced by semantic similarity. Gene Set Enrichment Analysis (GSEA) was performed on ranked gene lists based on shrunken log_2_ fold changes, using GO Biological Process gene sets and the same multiple-testing correction.

Targeted metabolic pathway enrichment was performed using curated *Arabidopsis* gene sets representing major metabolic and stress-related pathways, including photosynthesis/photorespiration, central carbon metabolism, the TCA cycle, amino acid and lysine metabolism, ROS/heat-stress responses, lipid metabolism, signaling and epigenetic regulation. Enrichment was evaluated using a hypergeometric test with all genes retained after DESeq2 filtering as the background.

Transcription-factor analyses were performed using a curated *Arabidopsis* TF database. TFs showing significant differential expression in at least one comparison were evaluated across basal genotype differences, heat and recovery responses, and ΔLFC values to identify genotype-dependent regulatory patterns. Gene annotation was performed using org.At.tair.db, and complete package versions and session information are provided in Supplementary Data.

## 3. Results

### DAPAT deficiency alters basal PSII efficiency but not recovery from prolonged warming

Reported elevations in free salicylic acid (SA) levels in *dapat* plants (Rate & Greenberg, 2001; Song *et al*., 2004), together with the metabolic reprogramming associated with the DAPAT mutation and the altered stress responsiveness of the mutant (Cavalcanti *et al*., 2018; Neves *et al*., 2024; Gouveia *et al*., 2025, *bioRxiv preprint*; Gouveia *et al*., 2026) (Supplementary Fig. S1b), prompted us to investigate how lysine deficiency influences plant responses to heat stress. Given the well-established role of SA in enhancing thermotolerance (Larkindale & Knight, 2002; Clarke *et al*., 2004), we hypothesized that the *dapat* mutant would show a distinct acclimation capacity under elevated temperature conditions. Wild-type (WT) and *dapat* plants were grown for three weeks under short-day (SD) conditions at 20 °C and then subjected to a sustained warming treatment at 26 °C (6 °C above control) for seven days, followed by a seven- day recovery period at 20 °C to assess phenotypic restoration. In parallel, a separate set of WT and *dapat* plants were maintained under control conditions for four weeks and then exposed to an acute heat shock (HS) treatment (38 °C for 6 h) (Supplementary Fig. S1c).

Phenotypic assessment showed that prolonged warming (PW) did not impair the recovery capacity of *dapat* plants (Figure 1a,b). To evaluate thermotolerance-associated markers related to photosystem integrity, the *maximum quantum efficiency of photosystem II* (*Fv/Fm*) was measured during warming and recovery (Supplementary Fig. S2a,b). *dapat* plants displayed lower *Fv/Fm* values even under control conditions, suggesting reduced PSII efficiency. Notably, PW further decreased *Fv/Fm* in WT plants but had no additional effect in the mutant. Together, these results indicate that, although *dapat* plants have intrinsically lower PSII efficiency, they maintain PSII performance under PW, whereas WT plants exhibit a further decline in *Fv/Fm* under the same conditions. Despite the growth inhibition caused by the DAPAT mutation, both WT and *dapat* plants displayed leaf epinasty in response to HS (Figure 1c). Because of the dynamic nature of the treatment, *Fv/Fm* was not measured under this condition.

**Figure 1.**
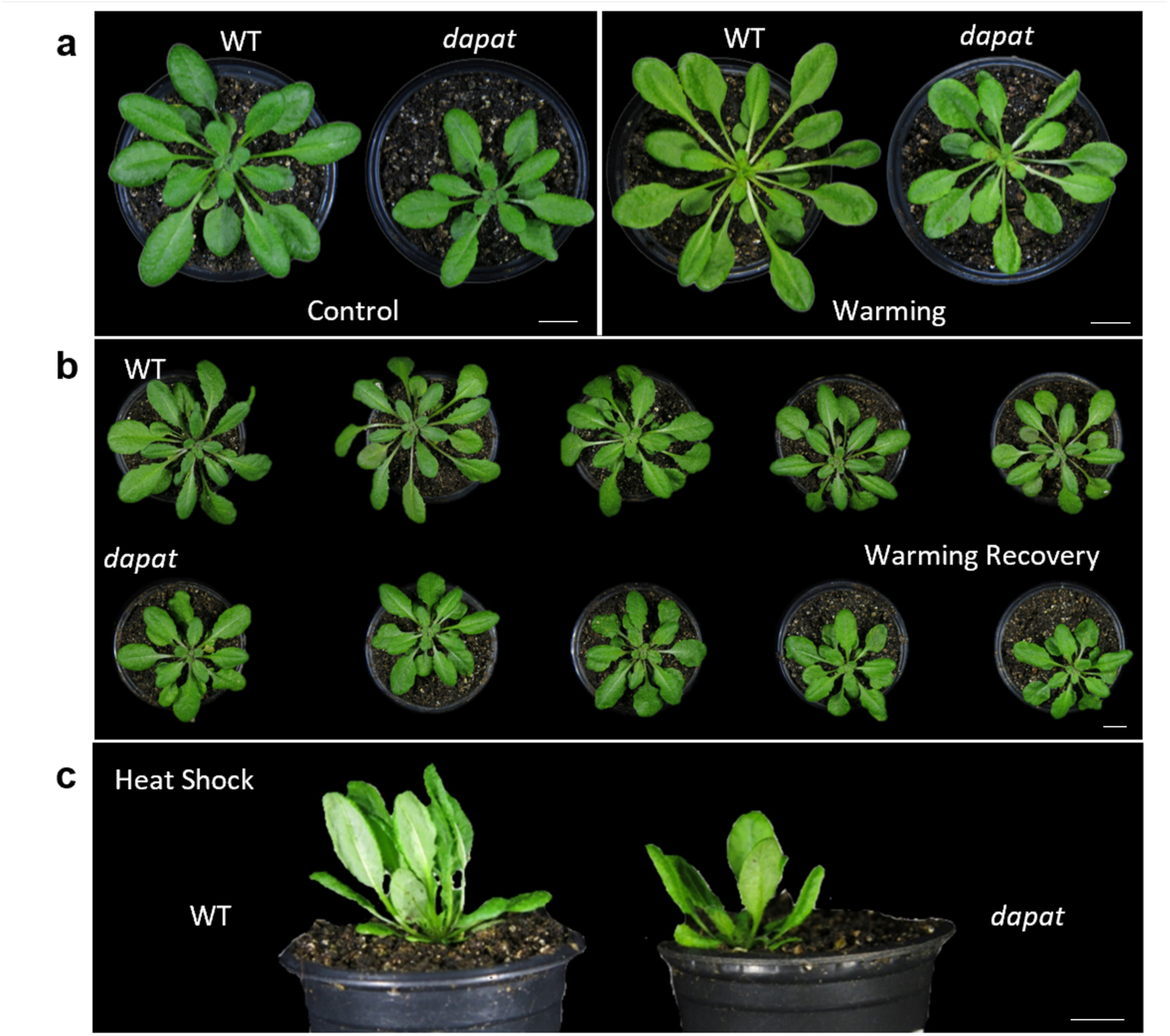
Phenotypic responses of wild-type and *dapat* plants to prolonged warming, heat shock and recovery. (a–c) Representative phenotypes of 5-week-old wild-type (WT) and *dapat* plants grown under control conditions (a), subjected to 7 days of prolonged warming (PW) (b), or exposed to heat shock (HS) followed by recovery (c). Plants were grown in pots under short-day conditions (8 h light/16 h dark) at 20 °C, with a light intensity of 150 µmol m□² s□¹ and 60% relative humidity. Plants were grown for 3 weeks before PW treatment or for 4 weeks before HS. The experimental workflow is shown in Fig. S1. **Abbreviations:** WT, wild-type; *dapat*, *dapat* mutant line.

### Lysine deficiency reshapes heat stress transcriptional responses in dapat mutant

As previously reported, *DAPAT* gene expression (*also named*, At4g33680), did not differ substantially between WT and *dapat* plants under control conditions (Cavalcanti *et al*., 2018; Gouveia *et al*., 2026). To examine whether *DAPAT* is differentially regulated by heat stress conditions, we quantified its expression in WT plants subjected to PW (Supplementary Fig. S2c) and in WT and *dapat* plants exposed to HS (Supplementary Fig. S2d). PW induced *DAPAT* expression in WT, reaching approximately 1.7-fold the control level (Supplementary Fig. S2c). Notably, HS strongly induced *DAPAT* expression in mutant plants, reaching approximately 2.0-fold the level observed under control, whereas no significant induction was detected in WT (Supplementary Fig. S2d). Thus, although *DAPAT* showed a stronger transcriptional response to HS in *dapat* than in WT plants, PW induced *DAPAT* expression in WT, whereas HS treatment did not significantly affect its expression. Because *DAPAT* expression was not evaluated in mutant plants under PW, these results suggest that, in WT, *DAPAT* responds preferentially to sustained rather than acute heat stress.

To further understand the molecular responses of WT and *dapat* plants to heat stress, we next examined the expression of genes associated with heat-stress perception and activation of the heat stress response (HSR). Targets were selected based on their strongest responses under PW and HS, which were analyzed independently. A core set of genes, including *HSFA1a, HSFA1b, HSFA1d, HSP90, HSP101, DREB2A, bZIP28, bZIP60, ANAC55, MBF1c, WRKY26*, and *WRKY33*, emerged as prominent components of this response (Figure 2). PW upregulated multiple heat-responsive genes in both genotypes (Figure 2a). Among the heat-shock proteins, *HSP90* transcript abundance was approximately 2.7-fold higher in *dapat* than in WT under control conditions; however, this difference was not maintained under PW. Moreover, *HSP101* was induced by PW, with no significant genotype-dependent differences. We next examined transcription factors from the HSF, DREB, WRKY, and bZIP families. *DREB2A*, *bZIP60*, and *WRKY26* were strongly induced by PW in both genotypes. *bZIP60* increased approximately threefold in WT and *dapat*, whereas the magnitude of *DREB2A* and *WRKY26* induction differed between genotypes. *DREB2A* expression increased approximately fivefold in WT and fourfold in *dapat* relative to WT control plants. By contrast, *WRKY26* increased approximately 2.5-fold in WT but approximately fivefold in *dapat*, indicating a markedly stronger response in the mutant. *WRKY33* expression was not significantly altered by PW in either genotype. Notably, *HSFA1a*, a master regulator of the HSR (Yoshida *et al*., 2011; Ohama *et al*., 2016, 2017), was preferentially induced in *dapat*, reaching approximately threefold higher transcript levels than in WT under PW.

**Figure 2.**
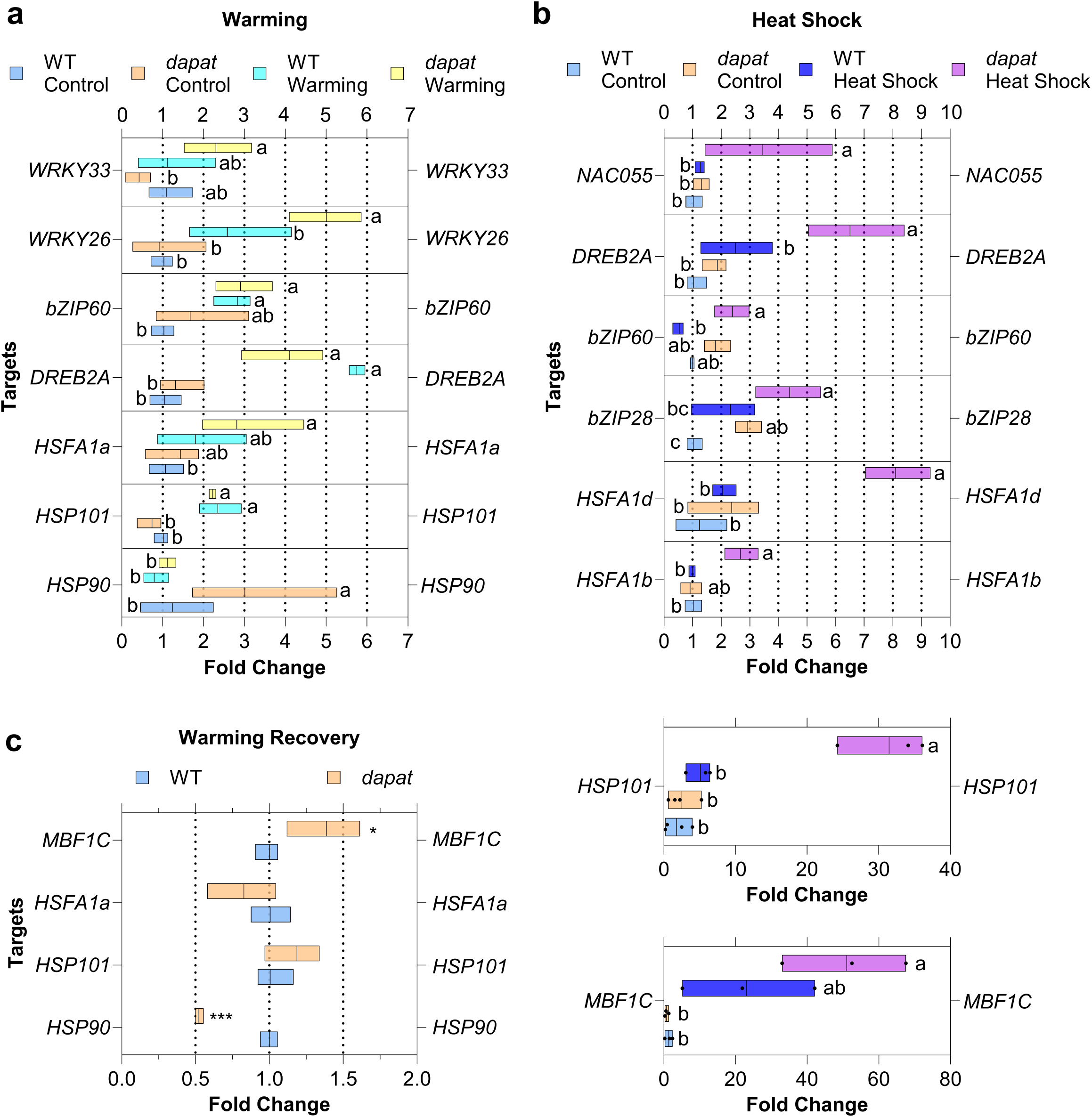
Differential regulation of heat-stress-responsive genes in wild-type and *dapat* plants. (a–c) Relative transcript levels of heat-stress-responsive genes in wild- type (WT) and *dapat* plants subjected to prolonged warming (PW) (a), heat shock (HS) (b), or recovery following PW (c) Box plots show transcript fold changes for the indicated transcription factors and heat-shock proteins: *WRKY33*, *WRKY26*, *bZIP60*, *DREB2A*, *HSFA1a*, *HSFA1d*, *HSFA1b*, *HSP101*, *HSP90* and *NAC055*. Panel c also shows transcript levels during recovery together with the corresponding HS responses of HSP101 and MBF1c. Center lines indicate means, and box limits indicate the 10th– 90th percentiles. Different lowercase letters indicate significant differences among treatments and genotypes, as determined by two-way ANOVA followed by Tukey’s multiple-comparisons test (P < 0.05). Asterisks indicate significant differences from the WT control (P < 0.05 or P < 0.001; two-tailed Student’s *t*-test). **Abbreviations:** WT, wild-type; *dapat*, *dapat* mutant line; bZIP, basic leucine zipper; DREB, dehydration- responsive element-binding protein; HSFA, heat-shock factor A; HSP, heat-shock protein; MBF1c, multiprotein bridging factor 1c; NAC, NAM, ATAF and CUC domain-containing transcription factor; WRKY, WRKY domain-containing transcription factor.

To determine whether these transcriptional differences persisted after stress, we quantified selected stress-responsive genes following a 7-day recovery period (Figure 2c). We also included *MBF1c*, a transcriptional coactivator previously associated with the establishment of thermotolerance in *Arabidopsis* (Suzuki *et al*., 2008). *HSFA1a* and *HSP101* transcript levels did not differ significantly between genotypes after recovery. In contrast, the elevated basal abundance of *HSP90* observed in *dapat* before stress (Figure 2a) was no longer maintained; during recovery, *HSP90* expression in *dapat* was approximately half that of WT. Conversely, *MBF1c* was upregulated in *dapat* after recovery, reaching approximately 1.4-fold higher levels than in WT (Figure 2c).

To extend this analysis, we examined the transcriptional response to HS conditions (Figure 2b). In contrast to the responses observed under PW, HS revealed pronounced genotype-dependent differences. Based on the response to acute high-temperature exposure, we examined *HSFA1b*, *HSFA1d*, *HSP90*, *DREB2A*, *bZIP28*, *bZIP60*, and *ANAC055* as targets. Most of these genes (*e.g. HSFA1b*, *HSFA1d*, *HSP90*, *bZIP28*, and *DREB2A*), were more strongly induced in *dapat*, with increases of approximately 2.0-, 3.0-, 2.0-, 3.0-, and 3.0-fold, respectively. By contrast, *bZIP60* expression did not differ significantly between genotypes after HS exposure.

### SA biosynthesis and signaling mutants reveal a distinct heat-stress transcriptional profile in *dapat*

To determine whether elevated endogenous SA contributes to the HSR phenotype of the *dapat* mutant, we included the SA biosynthesis mutant *sid2-1* (Wildermuth *et al*., 2001) and the SA signaling mutant *npr1-3* (Cao *et al*., 1997) as experimental controls (Figure 3). As in the first part of this study, WT, *dapat*, *sid2-1*, and *npr1-3* plants were germinated and grown under control conditions before being exposed to PW and HS, with the latter, now followed by a 24-h recovery period.

**Figure 3.**
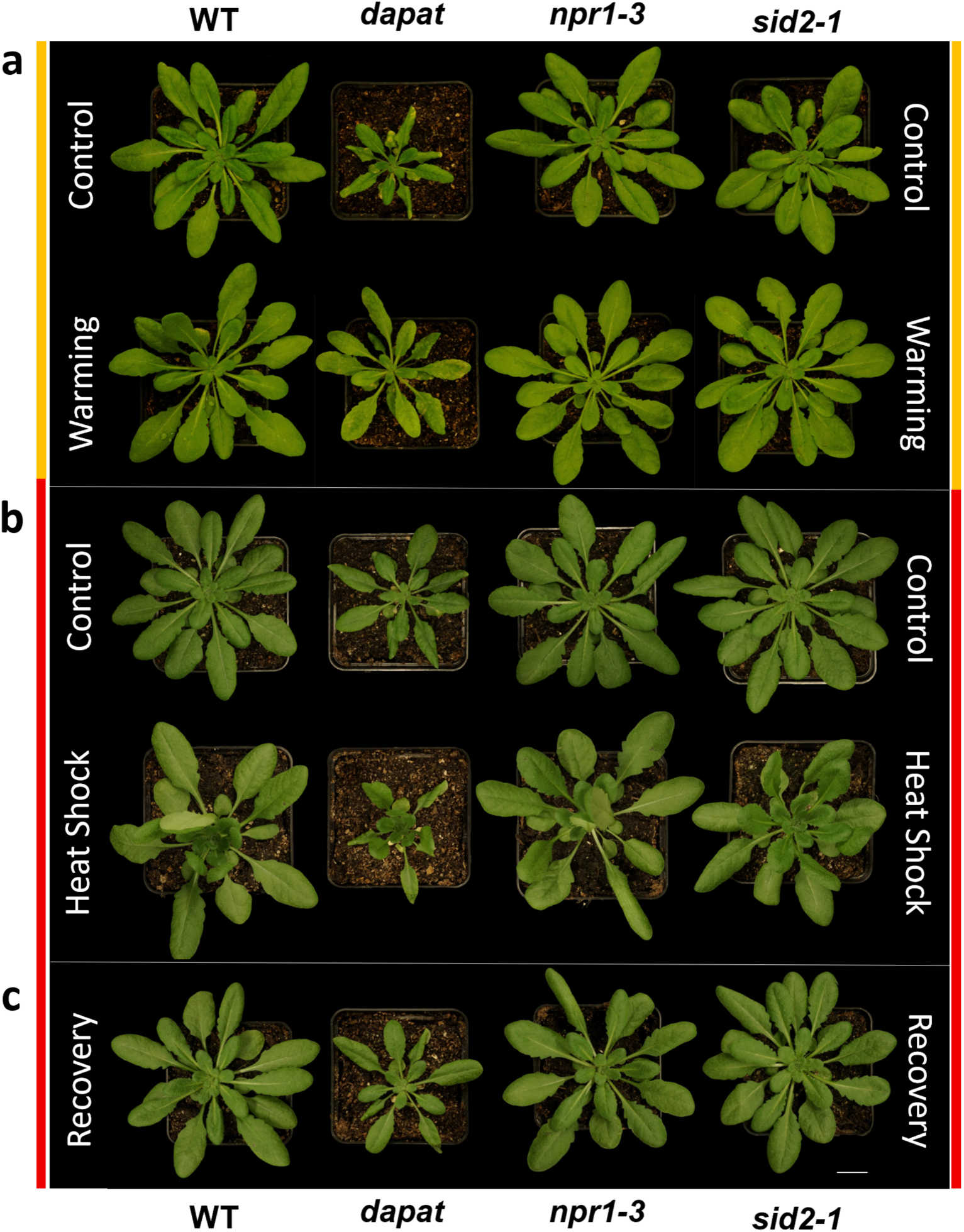
Phenotypic responses of wild-type, *dapat*, *npr1-3* and *sid2-1* plants to heat stress and recovery. (a–c) Representative phenotypes of 5-week-old wild-type (WT), *dapat*, *npr1-3* and *sid2-1* plants grown under control conditions (a), subjected to prolonged warming (PW) (b), or exposed to heat shock (HS) followed by recovery (c). Plants were grown under neutral-day conditions (12 h light/12 h dark) at 22 °C during the day and 20 °C at night, with a light intensity of 150 µmol m□² s□¹ and 60% relative humidity. **Abbreviations:** WT, wild-type; *dapat*, *dapat* mutant line; *npr1-3*, *nonexpressor of pathogenesis-related genes 1-3*; *sid2-1*, *salicylic acid induction deficient 2-1*.

Transcript levels of HSR genes were assessed after HS exposure, as this represents the most acute phase of stress imposition. Accordingly, we quantified the transcript levels of *HSFA1a*, *bZIP60*, *HSP90*, and *DREB2A* (Figure 4a). Under control conditions, *dapat* plants showed elevated *HSFA1a* and *bZIP60* transcript levels, consistent with a pre-activated stress state. Following HS, *HSFA1a* expression remained unchanged in *dapat* but was strongly induced in *sid2-1* and *npr1-3*. Likewise, *bZIP60* induction was weaker in *dapat* than in the SA-pathway mutants. *HSP90* was induced in all mutant backgrounds, with the strongest response in *npr1-3*, followed by *sid2-1* and *dapat*. Basal *DREB2A* expression was comparable in WT, *dapat*, and *sid2-1*, but slightly lower in *npr1-3*. After HS, *DREB2A* transcript abundance declined in all mutants, reaching its lowest level in *dapat* (Figure 4a).

**Figure 4.**
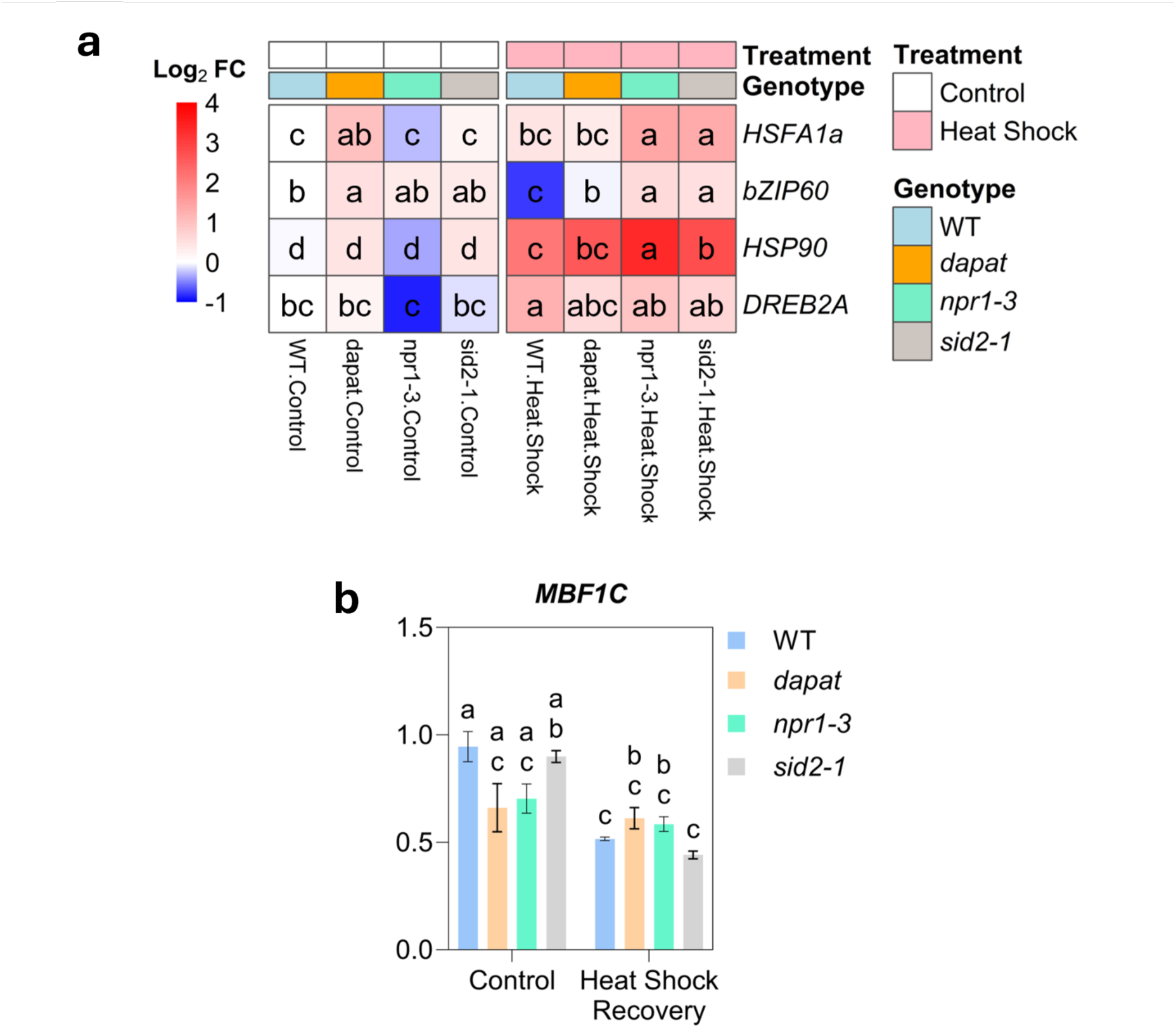
Regulation of heat-stress and unfolded-protein-response marker genes in wild-type and mutant plants. (a–b) Heat map showing log fold changes (log FC) of *HSFA1a*, *bZIP60*, *HSP90* and *DREB2A* transcript levels in wild-type (WT), *dapat*, *npr1-3* and *sid2-1* plants under control and after heat shock (a), relative to the corresponding control; colour intensity represents log FC values from −1 (blue) to 4 (red). Relative transcript levels of *MBF1c* (b) under control conditions and after post-heat-shock recovery. Data in a, b, d and e are means ± s.e.m. Different lowercase letters indicate significant differences among genotypes and treatments, as determined by two-way ANOVA followed by Tukey’s multiple-comparisons test (P < 0.05). **Abbreviations:** WT, wild type; *dapat*, *dapat* mutant line; *npr1-3*, *nonexpressor of pathogenesis-related genes 1-3*; *sid2-1*, *salicylic acid induction deficient 2-1*; HSFA1, heat-shock factor A1; bZIP60, basic leucine zipper transcription factor 60; HSP90, heat-shock protein 90; DREB2A, dehydration-responsive element-binding protein 2A; MBF1c, multiprotein bridging factor 1c; s.e.m., standard error of the mean.

During the 24-h recovery phase, *MBF1c* was assessed as a thermotolerance-associated marker (Figure 4b). Under control conditions, mutant lines showed reduced *MBF1c* expression, with the lowest levels in *dapat* and *sid2-1*. After HS and recovery, *dapat* and *sid2-1* showed elevated *MBF1c* expression relative to their controls, whereas *npr1-3* displayed a slight reduction. Together, these results show that *dapat* plants display a distinct heat stress transcriptional response compared with WT, with stronger activation of key HSR regulators under HS, with altered responses that persist into the recovery phase.

### Heat stress reveals distinct gas-exchange responses in *dapat* and SA-signaling mutants

Given the genotype-specific responses observed for heat-responsive gene expression, parameters in WT, *dapat*, *sid2-1*, and *npr1-3* plants following heat-stress treatment. Measurements included stomatal conductance (*g* apparent transpiration rate (*E*), leaf vapor pressure deficit (*VPD_leaf_*), photosystem II quantum efficiency (Φ*PSII*), and electron transport rate (*ETR*). These measurements revealed genotype-dependent physiological responses to PW and to HS followed by recovery (Table 1). Overall, gas-exchange parameters were more strongly affected than photosynthetic parameters.

The mutant lines differed physiologically from WT under control and PW conditions. Under control conditions, *gs* and *E* were broadly comparable across genotypes, with *dapat* exhibiting the lowest basal *gs* (0.182 µmol m□²s□¹). As expected, PW caused a general reduction in *E* across genotypes (–33% WT, –28% *dapat*, –34% *sid2-1* and –24% *npr1-3*,) consistent with stomatal closure under increased atmospheric demand, as reflected by a slight increase in *VPD_leaf_* to WT and SA- mutants. Φ*PSII* remained largely stable under warming, supporting PSII stability during PW induction. Strikingly, *ETR* increased by 34% in WT (2.713 to 3.642 µmol m□² s□¹), by 76% in *sid2-1* (2.057 to 3.678 µmol m□² s□¹), and by 76% in *npr1-3* (1.921 to 3.369 µmol m□² s□¹), remaining stable in *dapat* (2.087 to 2.138 µmol m□² s□¹). Based in the higher pre-stress *gs* and *E* observed in *sid2-1* and *npr1-3*, *dapat* plants appear to employ a more conservative stomatal-regulation strategy, a trait that shapes their distinct response to warming (Table 1).

After HS followed by recovery, all genotypes showed a marked reduction in *gs*, with values converging to 0.178–0.208 mol m□² s□¹ in WT, *dapat*, *npr1-3*, and *sid2-1* and *E* (2.012–2.742 mmol m□² s□¹), indicating a consistent post-stress decrease in water loss. By contrast, *VPD_leaf_* varied little among genotypes, suggesting a partial decoupling between atmospheric demand and stomatal responses. Φ*PSII* values were also largely maintained during recovery (∼0.746–0.761), indicating preservation of core photochemical efficiency after HS. Notably, *ETR* was maintained or slightly enhanced in some genotypes, with *npr1-3* reaching 4.213 μmol m□² s□¹ and 3.365 μmol m□² s□¹ in *sid2-1*, consistent with a compensatory adjustment of photosynthetic electron transport during recovery (Table 1). In summary, Pw and HS therefore imposed distinct physiological constraints. PW affected transpiration and electron transport in a genotype-dependent manner, whereas HS followed by recovery caused a broadly similar reduction in water loss while preserving PSII efficiency. Compared with SA-related mutants, *dapat* plants differed mainly in the magnitude and coordination of stomatal and electron-transport responses, consistent with a role for DAPAT-associated metabolism in coupling heat-induced gas-exchange regulation to photosynthetic acclimation.

### Salicylic acid dynamics during prolonged warming in *dapat* and salicylic acid- mutants

Building in the interplay between SA, the HSR and thermotolerance establishment, we quantified free SA, salicylic acid glucoside (SAG), salicylic acid glucose ester (SGE), and total SA in WT, *dapat*, *sid2-1*, and *npr1-3* plants subjected to 7-days of PW (Figure 5a). Samples were harvested on days 0, 2, 4, and 7 of warming, with sampling performed at the middle of the light period. For all quantified metabolites, *dapat* plants exhibited the highest levels at day 0, prior to PW in 3-week-old plants, and maintained elevated levels throughout the stress period, followed by *npr1-3* and WT. Warming treatment caused a pronounced decline in free SA in all genotypes relative to their respective controls (Figure 5a). Notably, reductions in SAG and total SA were observed only in *dapat*. Moreover, SGE levels rose exclusively in *dapat* under warming, whereas *sid2-1* and *npr1-3* remained unchanged. These findings underscore genotype specific metabolic adjustments to elevated temperature, with *dapat* exhibiting the most marked alterations (Figure 5a).

**Figure 5.**
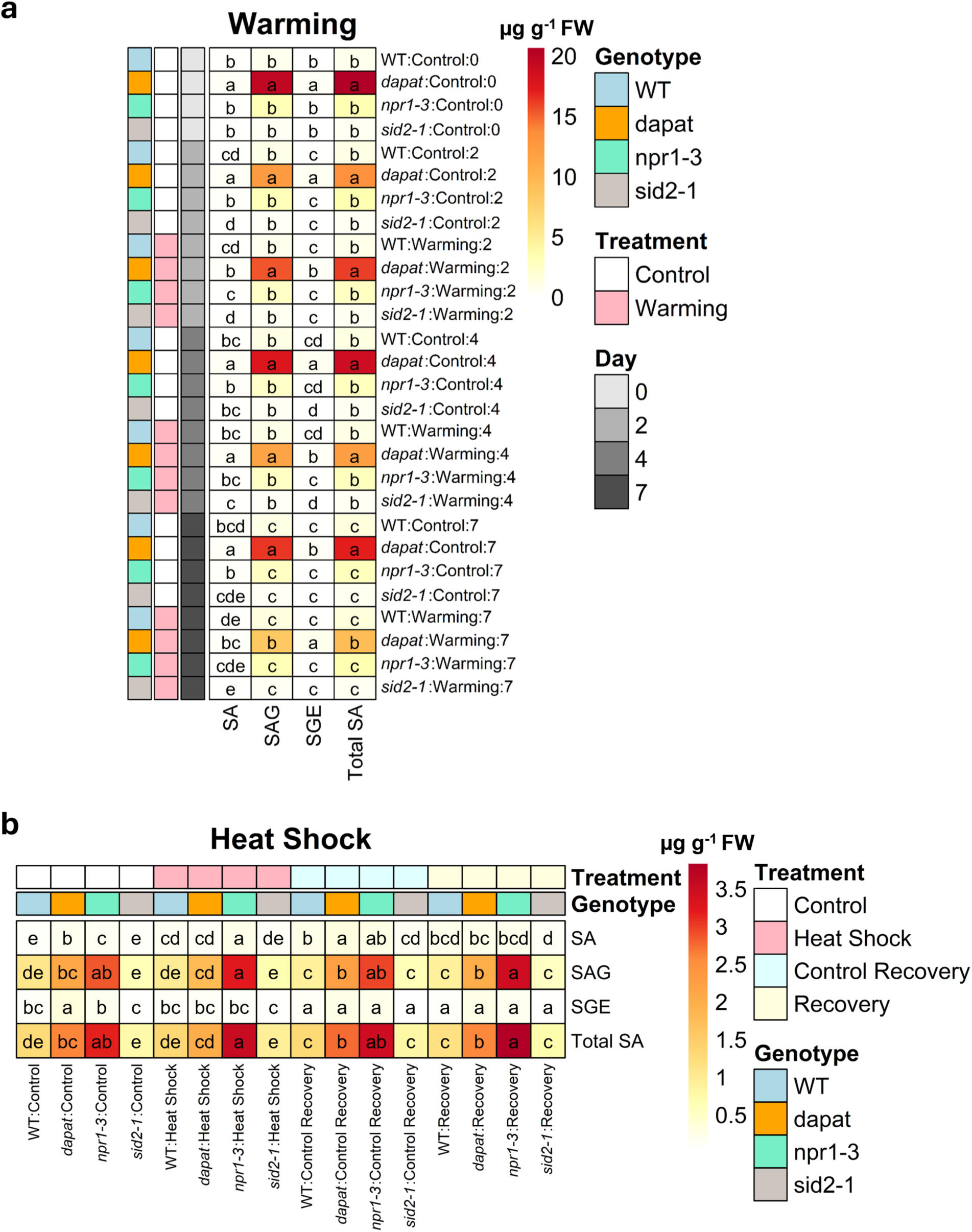
Salicylic acid dynamics during heat stress and recovery. (a,b) Heat maps showing the absolute abundance of free salicylic acid (SA), salicylic acid 2-O-β-D-glucoside (SAG), salicylic acid glucose ester (SGE) and total SA in wild-type (WT), *dapat*, *npr1-3* and *sid2-1* plants. (a) Metabolite levels under control conditions, heat shock (HS), control recovery and post-HS recovery. (b) Metabolite levels during a 7-day time course of prolonged warming, measured on days 0, 2, 4 and 7 under control and warming conditions. Concentrations are expressed as µg g ¹ fresh weight (FW); colour scales range from 0 to 3.5 µg g ¹ FW in a and from 0 to 20 µg g ¹ FW in b. Total SA represents the sum of free SA, SAG and SGE. Different lowercase letters indicate significant differences among genotypes within each treatment in a, or among genotypes at each time point in b, as determined by two-way ANOVA followed by Tukey’s multiple-comparisons test (P < 0.05). **Abbreviations:** WT, wild-type; *dapat*, *dapat* mutant line; *npr1-3*, *nonexpressor of pathogenesis-related genes 1-3*; *sid2-1*, *salicylic acid induction deficient 2-1*; SA, free salicylic acid; SAG, salicylic acid 2-O-β-D-glucoside; SGE, salicylic acid glucose ester; FW, fresh weight.

The next inquiry was to evaluate free SA, SAG, SGE, and total SA in WT, *dapat*, *npr1-3*, and *sid2-1* lines under HS and after 24h of post-stress recovery (Figure 5b). Samples were harvested for control and HS group (6h under 38°C), and after recovery period (24h), with sampling performed at the middle of the photoperiod. HS induced divergent SA dynamics across genotypes. Free SA levels increased in WT, *sid2- 1*, and *npr1-3*, whereas *dapat* exhibited a pronounced depletion, reaching WT- equivalent levels upon stress (Figure 5b). Notably, upon HS, *dapat* plants exhibited depletion of both SAG and SGE, with both pools returning to basal levels after recovery, whereas SAG and SGE levels in WT, *sid2-1*, and *npr1-3* remained comparable to control throughout stress and recovery. Finally, total SA was uniquely reduced in *dapat* by HS induction. After recovery, total SA levels reverted to control-like patterns, with *dapat* and *npr1-3* exhibiting the highest totals, followed by WT, and *sid2-1* the lowest (Figure 5b). Alternative comparison is provided in the supplementary material (Supplementary Figs. S3 and S4), which allows assessment of levels of free SA, SAG, SGE, and total SA (sum of SA, SA-O-glucoside [SAG], and SA glucose ester [SGE]) in *Arabidopsis* WT, *dapat*, *sid2-1* and *npr1-3* plants under PW and HS followed by recovery phase.

Taken together, these results show that *dapat* undergoes the most pronounced and genotype specific reconfiguration of SA pools under both PW and HS, with dynamic shifts in free and conjugated SA that contrast with the more stable profiles of WT, *sid2 1*, and *npr1 3*.

### DAPAT deficiency establishes a distinctive metabolic state during heat stress

The analysis of the metabolism dynamics in WT and *dapat*, *sid2-1*, and *npr1-3* plants subjected to PW and sampled throughout the stress (middle of the light period, MD; days 0, 2, 4, and 7) enabled us to assess the individual responses of each mutation and their behavior under treatment. Each cell represents the row-wise Z-scores, with red indicating higher and blue lower metabolite abundance relative to the row mean (Figure 6). Prior to stress, *dapat* display a distinct metabolic signature compared to SA–related mutants. MD sampling revealed a pronounced broad accumulation of amino acids, alongside selective depletion of proline. Notably, intermediates of the TCA citrate: isocitrate levels and organic acids (*e.g*., fumaric, isocitric, maleic, malic, oxoglutaric, pyruvic and succinic) also accumulated substantially. In terms of carbohydrates, sucrose levels increased while reducing sugars (glucose and fructose) were depleted (Figure 6). Amino acid metabolism exhibits the most complex and genotype-dependent responses to PW. In summary, WT significant depleted several amino acids (*e.g*., glycine, serine, threonine, glutamine, asparagine, aspartate, phenylalanine, and proline), with the latter being strongly depleted across genotypes by day 7. Conversely, *dapat* sustained the amino acid enrichment observed under control, including GABA, lysine, and BCAA (leucine, isoleucine, and valine). This profile contrasts with the *sid2-1* and *npr1-3*, which shared only lysine accumulation alongside proline depletion (Figure 6).

**Figure 6.**
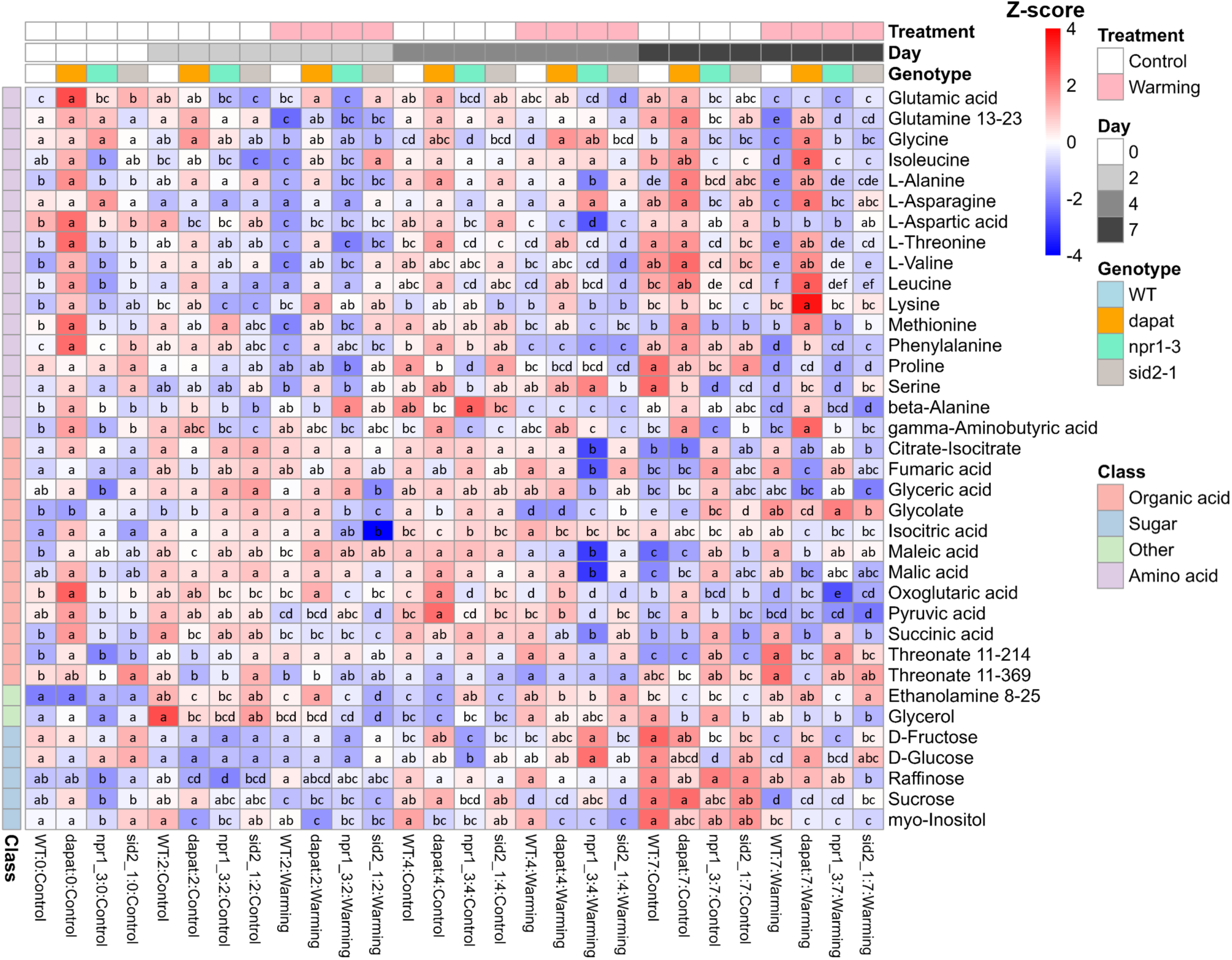
Temporal metabolic profiling of wild-type and mutant plants during prolonged warming. Relative abundance of primary metabolites in wild-type (WT), *dapat*, *npr1-3* and *sid2-1* plants under control and prolonged-warming conditions over a 7-day time course, with measurements collected on days 0, 2, 4 and 7. Values are expressed as Z-scores calculated across row means, ranging from −4 (blue, lower abundance) to 4 (red, higher abundance). Different lowercase letters within individual heat-map cells indicate significant differences among genotypes at each time point and treatment, as determined by two-way ANOVA followed by Tukey’s multiple-comparisons test (P < 0.05). Metabolites are grouped by biochemical class—amino acids, organic acids, sugars and other metabolites, as indicated by the colour key. Numeric suffixes adjacent to metabolite names (for example, *Ethanolamine 8–25*) denote chromatographic retention times in minutes. **Abbreviations:** WT, wild-type; *dapat*, *dapat* mutant line; *npr1-3*, *nonexpressor of pathogenesis-related genes 1-3*; *sid2- 1*, *salicylic acid induction deficient 2-1*; Z-score, standardized deviation from the mean.

Additionally, organic acids exhibited pronounced modulation to PW across genotypes and time points. PW induced accumulation of isocitrate, maleate, succinate and glycolate, whereas 2-oxoglutarate and pyruvate were markedly depleted in all genotypes, particularly at the end of the treatment (Figure 6). The sugar profile revealed a trend toward depletion of soluble sugars in WT and *npr1-3*. Equally noteworthy was the sustained accumulation of raffinose in *dapat* and *npr1-3* under both control and PW by day 7 (Figure 6). Together, these results show that PW induces a dynamic and genotype-dependent metabolic response. Although all genotypes displayed substantial changes in amino-acid, organic-acid, and sugar metabolism, *dapat* maintained a distinct amino-acid-enriched profile throughout the treatment, which was not fully reproduced by the SA-pathway mutants. We next asked whether this metabolic signature was also evident under acute heat stress and during the subsequent recovery phase.

To address this question, we profiled metabolites in WT, *dapat*, *sid2-1*, and *npr1-3* plants exposed to HS (38°C for 6h) after 24h recovery, with samples collected at midday. As observed under PW, *dapat* showed the most distinct basal metabolome relative to WT and the SA-pathway mutants. Under control conditions, *dapat* exhibited elevated levels of multiple amino acids, including β-alanine, glycine, alanine, threonine, valine, methionine, glutamine, and proline, as well as increased organic acid intermediates (*e.g*., citrate: isocitrate levels, fumaric acid, malic acid, and glutaric acid). In contrast, raffinose was markedly depleted. The SA-pathway mutants displayed more similar basal profiles, characterized by reduced glutamate, glycine, serine, glucose, and fructose, alongside increased citrate:isocitrate, fumarate, and malate; raffinose accumulation was detected only in *sid2-1* (Figure 7).

**Figure 7.**
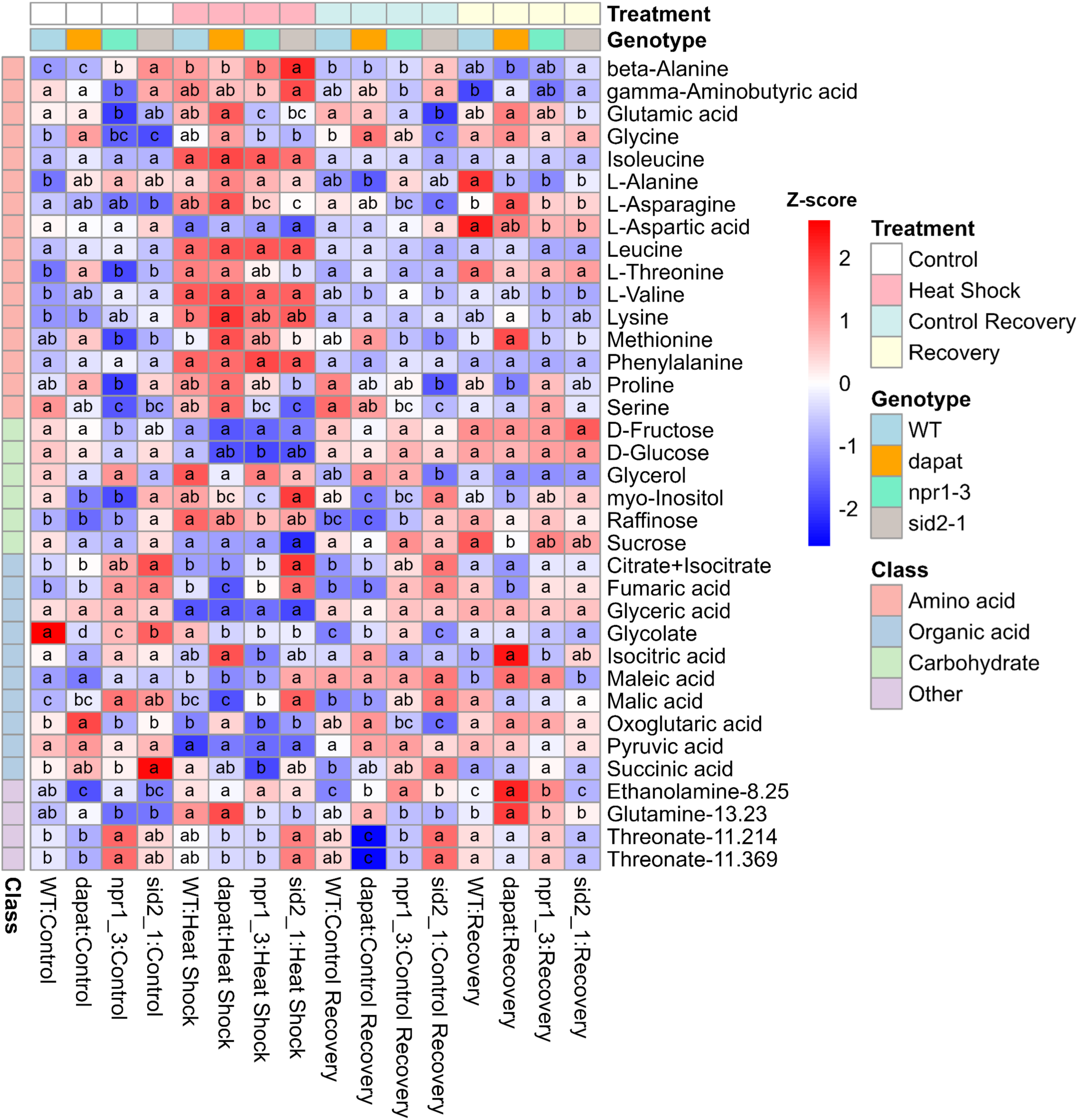
Metabolic profiles of wild-type and mutant plants during heat stress and recovery. Relative abundances of primary metabolites in wild-type (WT), *dapat*, *npr1-3* and *sid2-1* plants under control conditions, heat shock (HS), control recovery and post- HS recovery. Values are shown as row-wise Z-scores, with red indicating higher and blue lower metabolite abundance relative to the row mean. Different lowercase letters within individual cells indicate significant differences among genotypes within each treatment, as determined by two-way ANOVA followed by Tukey’s multiple- comparisons test (P < 0.05). Metabolites are grouped by biochemical class, as indicated by the vertical colour key. Numbers following metabolite names (for example, *ethanolamine-8.25*) denote chromatographic retention times in minutes. **Abbreviations:** WT, wild type; *dapat*, *dapat* mutant line; *npr1-3*, *nonexpressor of pathogenesis-related genes 1-3*; *sid2-1*, *salicylic acid induction deficient 2-1*; Z-score, standardized value relative to the row mean.

Following HS, all genotypes showed broad amino-acid accumulation, particularly of β-alanine, BCAA, and lysine. In contrast, glucose, fructose, glycerate, and pyruvate decreased, whereas raffinose accumulated strongly across all genotypes (Figure 7). Thus, in contrast to the more genotype-dependent response to PW, acute HS elicited a more convergent metabolic response, with increased amino acids and raffinose and reduced soluble sugars and central-carbon intermediates. Upon recovery, lysine and BCAA declined across genotypes, while glucose and fructose increased, reversing their HS-induced depletion. Raffinose remained elevated, and TCA-cycle intermediates were partially restored; however, malate remained strongly depleted in WT and *sid2-1* (Figure 7). Notably, the recovery-phase metabolome of *dapat* resembled that of WT immediately after HS, although *dapat* uniquely accumulated asparagine and glutamine relative to WT during recovery.

Together, PW and HS reveal that DAPAT deficiency establishes a distinct basal metabolic state that persists during PW but becomes less evident after HS exposure. Both heat conditions reconfigured amino-acid, sugar, and organic-acid metabolism, whereas HS produced a broadly shared response across genotypes.

### Principal component analysis reveals distinct metabolic reprogramming in *dapat* under heat stress

To integrate the metabolite-specific changes described above and better describe the overall relationships among genotypes, heat treatments, and sampling times, we performed principal component analysis (PCA). Under PW, PC1 and PC2 explained 32.3% and 18.6% of the total variance, respectively (Supplementary Fig. S5a). PC1 primarily separated control and warming samples and was driven by amino acids, particularly BCAA, threonine, and glutamine. PC2 mainly resolved samples according to treatment duration, with strong contributions from TCA-cycle and respiratory intermediates, including pyruvate, succinate, malate, citrate:isocitrate, and glycerate-3- phosphate. The PCA also revealed genotype-specific patterns during warming. *dapat* showed the largest PC1 shift, consistent with its marked accumulation of BCAA and broader metabolic reprogramming. By contrast, WT showed a smaller shift, whereas *sid2-1* and *npr1-3* clustered in intermediate positions, supporting genotype-dependent modulation of metabolic responses to PW.

Next, the PCA of HS and recovery datasets showed distinct changes. PC1 and PC2 explained 31.2% and 25.2% of the total variance, respectively, accounting for 56.4% of the overall variation (Supplementary Fig. S5b). PC1 separated HS samples according to the accumulation of amino acids, including leucine, valine, β-alanine, and phenylalanine, and the depletion of glycerate. PC2 mainly distinguished recovery samples, which were associated with changes in asparagine, glutamine, glutamate, and methionine, whereas citrate:isocitrate contributed negatively. Consistent with the univariate analysis, *dapat* showed the largest PC1 displacement after HS, reflecting the extensive amino-acid accumulation. WT exhibited a more moderate heat-induced shift, while *sid2-1* and *npr1-3* also displayed intermediate positions. Thus, PCA independently supports the conclusion that *DAPAT* deficiency amplifies metabolic reconfiguration during heat stress, whereas SA-pathway mutations modify, but do not mirror, the *dapat* metabolic phenotype.

### Genome-wide transcriptional reprogramming distinguishes *dapat* from WT during heat stress and recovery

To identify the transcriptional signatures underlying the divergent physiological and metabolic behavior of *dapat* under heat stress, we performed RNA sequencing in WT and *dapat* plants across four conditions: control-HS, HS, control-recovery, and HS- recovery. For similarity analyses, the data were VST-transformed (Supplementary Fig. S6a). The resulting sample-distance matrix showed consistent clustering of biological replicates and clear separation of HS samples from control and recovery conditions, supporting the reproducibility of the transcriptional profiles (Supplementary Fig. S6b). PCA of the VST-transformed data showed that PC1 and PC2 explained 60% and 15% of the total variance, respectively. PC1 predominantly captured the transcriptional shift associated with HS, separating HS samples from control and recovery conditions, whereas PC2 reflected genotype-dependent variation, distinguishing WT from *dapat* samples across treatments in the y axis scores (Supplementary Fig. S7).

To further resolve the transcriptional patterns underlying the separation observed in the PCA, we performed hierarchical clustering of the 100 most variable genes across all groups (Figure 8).The largest and most coherent cluster comprised classical HSR and protein-folding genes, including several small heat-shock proteins (*HSP18.2*, *HSP17.8*, *HSP17.6C*, *HSP17.6B*, *HSP17.4*, *HSP23.5*), chaperones and co-chaperones (*AtHSP70*, *AtHS83*, *AtFKBP65*, *DNAJ*, *Hop3*), the mitochondrial isoform *AtHSP23.6-MITO*, and the heat-stress transcription factor targets *AtMBF1C* and *HSA32*. These genes were strongly and coordinately induced under HS in both genotypes, reaching their greatest magnitude in *dapat* HS samples before progressively returning towards baseline during recovery, consistent with the canonical and rapidly reversible cytoplasmic HSR.

**Figure 8.**
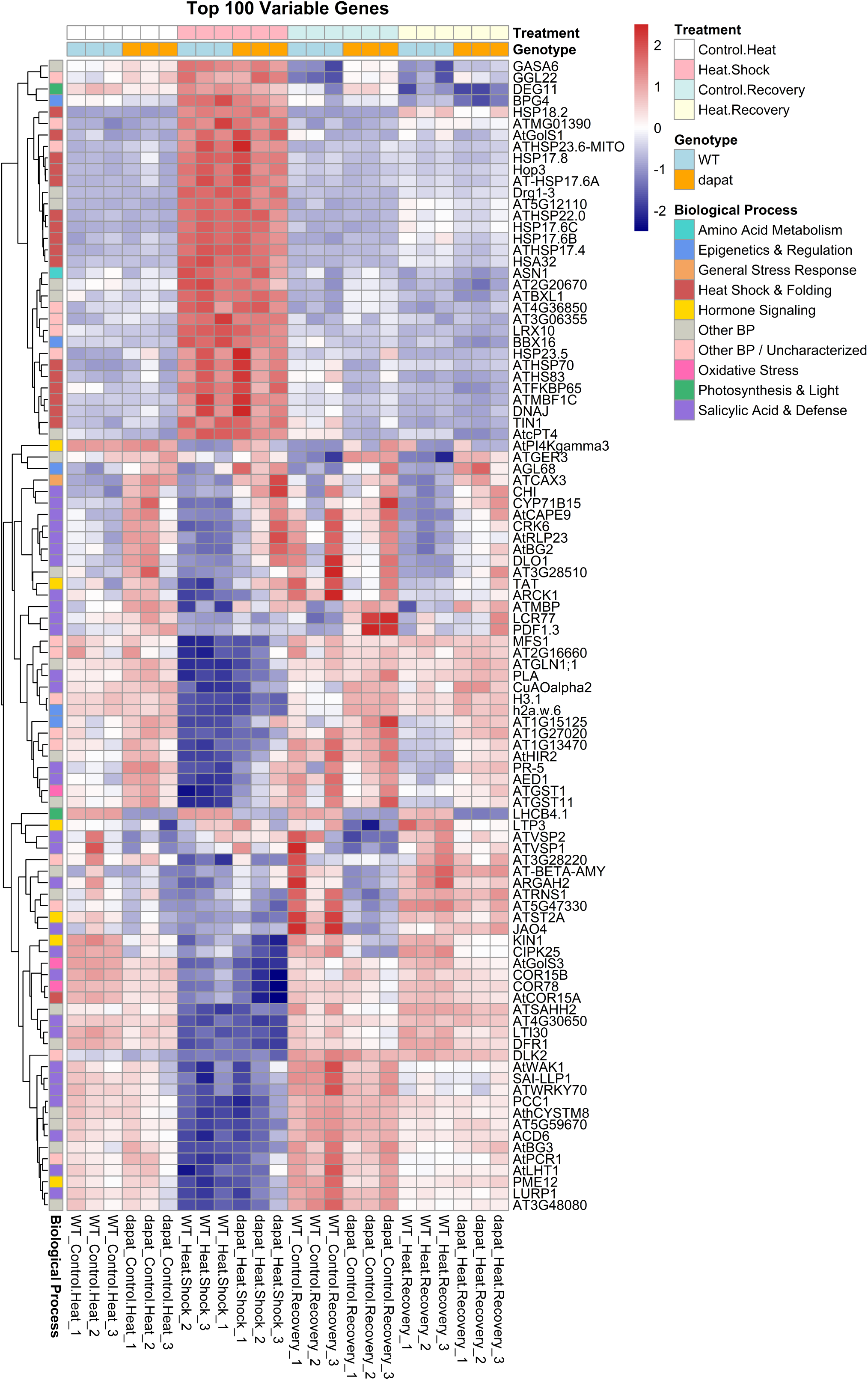
Transcriptomic signatures of wild-type and *dapat* plants during heat shock and recovery. Hierarchical-clustering heat map showing row-wise Z-scores for the 100 most variable genes identified by RNA sequencing in wild-type (WT) and *dapat* plants. The dataset comprises three independent biological replicates for each genotype under control, heat-shock, control-recovery and post-heat-shock-recovery conditions. Red indicates higher and blue lower transcript abundance relative to the mean expression of each gene; Z-scores range from −2 to 2. The vertical colour key denotes functional categories, including heat shock and protein folding, salicylic acid and defense, hormone signaling, general stress responses, photosynthesis and light responses, amino-acid metabolism, epigenetics and gene regulation, oxidative stress and other biological processes. **Abbreviations:** WT, wild-type; *dapat*, *dapat* mutant line; RNA-seq, RNA sequencing; Z-score, standardized value relative to the row mean.

A second major cluster, enriched for SA and defense-related genes (*PR-5*, *PDF1.3*, *AtWRKY70*, *PCC1*, *ACD6*, *AtBG3*, *AtHIR2*, *AED1*, *CRK6*, *AtRLP23*, *CYP71B15*) together with hormone-signaling genes (*TAT*, *ARCK1*, *KIN1*, *CIPK25*, *JAO4*), were generally repressed during HS and control-HS conditions but recovered, often to levels above control, during the recovery phase. Some of these transcripts (*e.g*., *PDF1.3*, *LCR77*) showed a distinctive peak specifically in *dapat* control-recovery samples, indicating a genotype-specific reactivation of defense-associated transcription independent of active heat stress. A distinct group comprising general stress markers (*COR15B*, *COR78*, *AtCOR15A*, *LTI30*, *KIN1*, *AtSAHH2*) and the photosynthesis/light- associated genes *LHCB4.1* and *GGL22* were strongly downregulated during HS and only partially restored during recovery, an effect more pronounced in WT than in *dapat*. This pattern parallels the sustained decline in Φ*_PSII_* -associated components (Table 1) and is consistent with photosynthetic gene repression as a shared feature of the acute heat response.

Specifically, amino-acid metabolism genes including *ASN1* (asparagine synthetase), *TAT* (tyrosine aminotransferase), *ARGAH2* (arginase), *At-BETA-AMY* (β- amylase), and *AtLHT1* (lysine/histidine transporter), together with epigenetic and chromatin regulators (*H3.1, h2a.w.6*) and oxidative-stress genes (*AtGST1*, *AtGST11*), formed smaller but distinct sub-clusters with variable induction patterns across HS and recovery, generally showing stronger modulation in *dapat* relative to WT, particularly during the recovery phase.

Functional enrichment analyses revealed a broadly conserved heat-stress response in both WT and *dapat* plants but also uncovered distinct genotype-dependent transcriptional programs and recovery trajectories. This suggests that the DAPAT mutation does not compromise canonical HSR but instead reshapes secondary transcriptional programs linked to defense signaling, photosynthesis, and amino-acid metabolism, consistent with a genotype-specific mode of transcriptional acclimation to heat stress. Under acute HS, GO enrichment analysis showed strong representation of processes associated with water deprivation, oxidative stress, and RNA modification in both genotypes (Supplementary Fig. S8a,d), indicating activation of core stress and homeostasis-related responses. Consistent with these findings, KEGG analysis identified carbon metabolism, cofactor biosynthesis, and starch and sucrose metabolism as major pathways responding to HS in both WT and *dapat* (Supplementary Fig. S8b,e). GSEA further supported this shared response, revealing coordinated changes in protein folding, thermal-response pathways, and reactive oxygen species regulation (Supplementary Fig. S8c,f). Together, these complementary analyses indicate that the loss of lysine biosynthetic capacity does not abolish the canonical transcriptional response to heat stress but rather alters the relative engagement of specific regulatory programs.

Genotype-dependent differences became more apparent when *dapat* was compared directly with WT. After HS exposure, *dapat* displayed stronger enrichment of defense and immune-related processes, together with signatures associated with SA signaling, oxidative stress, and glutathione metabolism (Supplementary Fig. S11a). KEGG analysis likewise highlighted plant–pathogen interactions, phenylpropanoid biosynthesis, and glutathione metabolism in *dapat* relative to WT (Supplementary Fig. S11b), while GSEA revealed coordinated shifts in SA and immune-associated gene sets (Supplementary Fig, S11c). Thus, although both genotypes activate a conserved heat- stress response, impaired lysine biosynthesis is associated with a greater engagement of defense- and redox-related transcriptional programs during acute heat stress.

These differences extended beyond the immediate stress response and became particularly pronounced during recovery. In WT plants, recovery was characterized by continued enrichment of defense-, immune-, and stress-responsive processes in GO analysis (Supplementary Fig. S9a), accompanied by strong representation of plant– pathogen interaction and protein processing in the endoplasmic reticulum in KEGG analysis (Supplementary Fig. S9b). GSEA similarly showed sustained enrichment of immune- and stress-associated gene sets (Supplementary Fig. S9c), indicating that WT plants maintain a stress-responsive state during the recovery phase. In contrast, *dapat* recovery samples highlighted a shift toward processes associated with cellular maintenance and restoration, including ribosome biogenesis, RNA modification, mitochondrial organization, and translational processes (Supplementary Fig. S9a,d). These features were further supported by KEGG enrichment of ribosome-related and RNA degradation pathways (Supplementary Fig. S9e) and by GSEA signatures associated with cytoplasmic translation, autophagy, and vacuole organization (Supplementary Fig. S9c,f). Collectively, these patterns indicate that *dapat* and WT follow distinct transcriptional pattern during recovery, with *dapat* preferentially engaging programs related to cellular reorganization and biosynthetic recovery rather than maintaining the predominantly defense-oriented state observed in WT.

Baseline genotype differences were also evident under control conditions, although they were less extensive than those observed following heat stress. Under control heat conditions, *dapat* versus WT comparisons were enriched for water deprivation, oxidative stress, SA signaling, and glutathione-associated processes (Supplementary Fig. S10a), with glutathione metabolism also emerging prominently in KEGG analysis (Supplementary Fig. S10b). GSEA further identified coordinated changes related to fatty acid catabolism and reactive oxygen species responses (Supplementary Fig. S10c), suggesting that impaired lysine biosynthesis is associated with altered redox and stress-related states even in the absence of acute heat shock. Under control recovery conditions, the genotype-dependent differences shifted toward DNA damage responses, ribosome biogenesis, and RNA-related processes in GO analysis (Supplementary Fig. S10d), with KEGG enrichment of ribosome biogenesis, DNA replication, and homologous recombination (Supplementary Fig. S10e) and GSEA signatures involving cell-cycle processes, RNA splicing, and jasmonic acid signaling (Supplementary Fig. S10f).

A pathway-level integration of transcriptomic and metabolomic data using Fisher’s combined-probability test (Supplementary Table S1) reinforced the main patterns above. Across all eight contrasts, metabolomic *P*-values were non-significant (*P* > 0.09), indicating that the joint pathway signals were driven predominantly by transcriptional changes. The shared heat-shock response included protein processing in the endoplasmic reticulum (WT, *P* = 2.0 × 10 ³; *dapat*, *P* = 3.0 × 10 ³), together with citrate-cycle, sugar and ascorbate/aldarate metabolism, consistent with coordinated proteostasis, carbon and redox responses. The *dapat*–WT divergence in defense and redox pathways was evident already under control conditions, with glutathione metabolism (*P* = 0.018) becoming markedly stronger under heat shock (*P* = 7.4 × 10□□), alongside plant–pathogen interaction (*P* = 3.9 × 10 ³) and phenylpropanoid biosynthesis (*P* = 1.6 × 10 ³), suggesting amplification of a constitutive defense state rather than de novo stress induction. The strongest signal in Supplementary Table S1 was a MapMan receptor-like kinase bin (875 genes) in *dapat* versus WT under HS (*P* = 9.9 × 10 ¹³), although its interpretation requires leading-edge analysis; this bin was also enriched in WT recovery versus control-recovery (*P* = 1.5 × 10□□).

The clearest genotype-specific recovery signature emerged from the direct *dapat*-versus-WT comparison at recovery period, where ribosome (*P* = 1.4 × 10□¹□), aminoacyl-tRNA biosynthesis (*P* = 2.7 × 10□□), starch and sucrose metabolism (*P* = 0.013) and lysine biosynthesis (*P* = 0.027; combined *P* = 0.017) were enriched. Lysine biosynthesis also showed the lowest metabolomic *P*-value in the dataset (0.092), suggesting the strongest, albeit non-significant, transcript–metabolite convergence. Consistently, *dapat* recovery samples versus control-recovery showed strong enrichment of ribosome (*P* = 3.6 × 10 ³) and ribosome biogenesis (*P* = 4.3 × 10□¹□), whereas neither pathway was enriched in WT. Additionally, ER protein processing remained prominent in WT recovery (*P* = 1.2 × 10□□) but was absent from *dapat*, supporting a transition from persistent proteostasis/defense in WT to anabolic recovery in *dapat*.

Together, these data support a model in which heat stress activates signaling pathways that converge on *HSFA1* and other heat-stress transcription factors, while SA biosynthesis and NPR1 signaling further modulate the core heat-stress response (Figure 9). In this model, genotype-specific differences in stomatal regulation, electron transport, and amino-acid and organic-acid metabolism act upstream of, and in parallel with, *HSFA1*/*DREB2A*-mediated transcriptional reprogramming, ultimately shaping the accumulation of stress-responsive genes that seems to determine the degree of stress tolerance achieved by each genotype. Within this framework, the DAPAT mutation appears to act primarily by shifting the amplitude and coordination of gas-exchange and metabolic responses rather than by disrupting the canonical *HSFA1*-centred transcriptional module, positioning DAPAT-associated lysine metabolism as a modulator that fine-tunes, rather than replaces, the SA- and NPR1-dependent branches of the HSR network.

**Figure 9.**
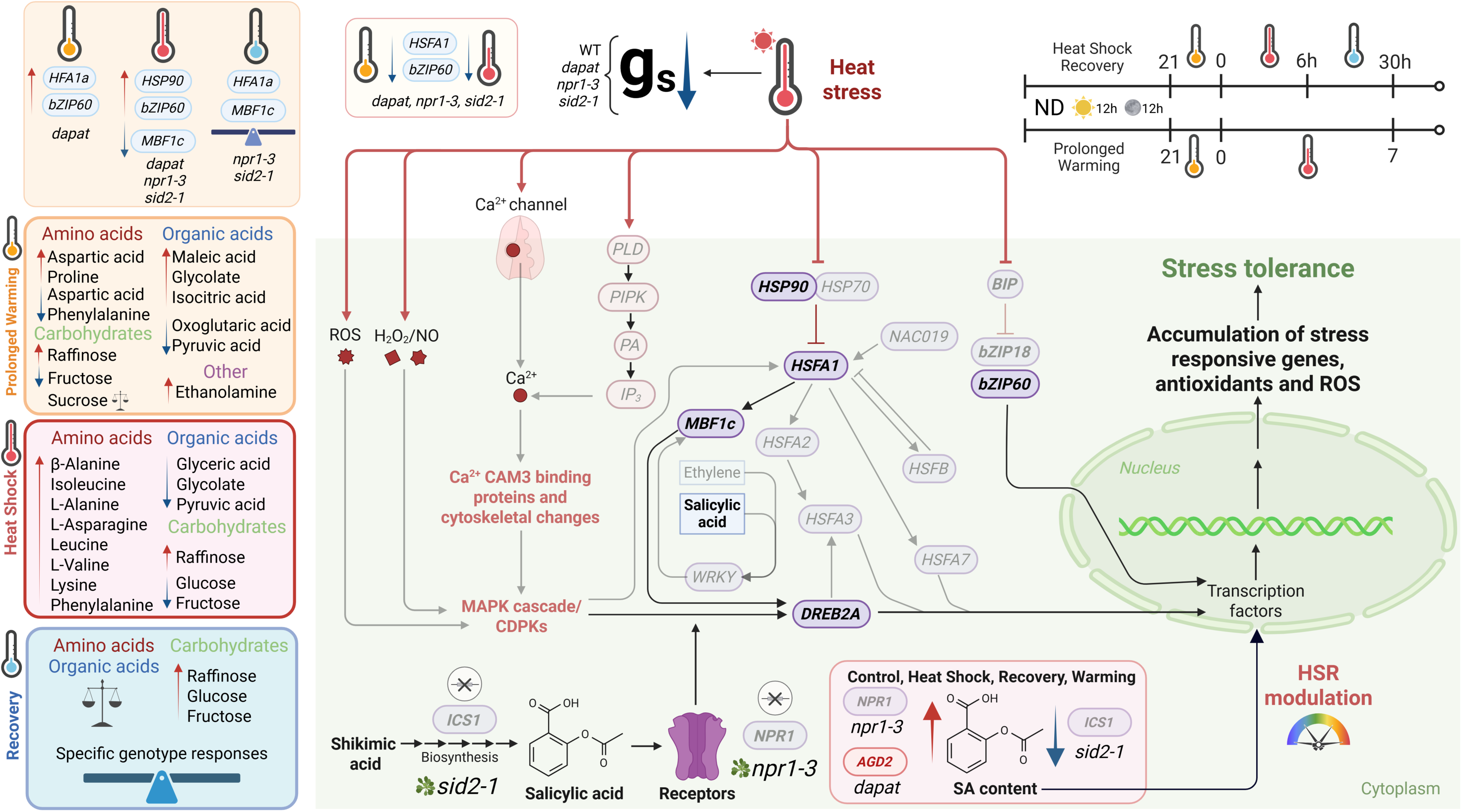
Integrative model of molecular, metabolic and hormonal responses to heat stress in *dapat mutant plants*. Schematic model summarizing the molecular, hormonal and metabolic responses of wild-type (WT), *dapat*, *npr1-3* and *sid2-1* plants to heat shock, recovery and prolonged warming. Plants were grown under neutral-day conditions (12 h light/12 h dark) at 22 °C before exposure to heat shock (38 °C for 6 h, followed by 24 h recovery at 22 °C during the day and 20 °C at night) or prolonged warming (6 °C above the control temperature for up to 7 days). Samples collected at 0, 6 and 30 h during the heat-shock experiment are indicated in the model. Canonical early heat-stress responses included the accumulation of reactive oxygen species (ROS), hydrogen peroxide (H O□□) and nitric oxide (NO), together with calcium influx through channels associated with the PLD–PIPK–PA–IP pathway. These signals are proposed to activate MAPK and CDPK cascades, CAM3-associated proteins and cytoskeletal remodelling. Heat stress also activated heat-responsive transcription factors, including *HSFA1*, *bZIP60*, *MBF1c* and *DREB2A*, as well as molecular chaperones such as *HSP90*, which promote the expression of downstream stress- tolerance genes. Heat stress reduced stomatal conductance (*gs*) across all genotypes, confirming effective stress imposition. SA biosynthesis through ICS1 and SA signaling through NPR1 were associated with the heat-stress response. The *sid2-1* and *npr1-3* mutants, impaired in SA biosynthesis and signaling, respectively, together with *dapat*, displayed distinct patterns of SA accumulation and heat-responsive gene expression under the different stress conditions. Prolonged warming was associated with the accumulation of selected amino acids, including Asp, Pro and Phe, organic acids, including Mal, Glyc and Iso, and raffinose (Raf). Heat shock induced broader amino- acid accumulation and carbohydrate depletion, followed by the restoration of carbohydrates, including Raf, Glc and Fru, during recovery. Opaque boxes indicate components directly evaluated in this study, whereas semi-transparent boxes indicate pathway intermediates inferred from previous literature. **Abbreviations:** AGD2, aberrant growth and development 2; bZIP, basic leucine zipper transcription factor; BiP, binding immunoglobulin protein; CAM3, calmodulin 3; CDPK, calcium-dependent protein kinase; *dapat*, *dapat* mutant line; DREB2A, dehydration-responsive element- binding protein 2A; Fru, fructose; Glc, glucose; Glyc, glycolate; `g_s`, stomatal conductance; HSFA/HSFB, heat-stress transcription factor A/B; HSP, heat-shock protein; HSR, heat-stress response; ICS1, isochorismate synthase 1; IP, inositol 1,4,5- trisphosphate; Iso, isocitric acid; Mal, maleic acid; MAPK, mitogen-activated protein kinase; MBF1c, multiprotein bridging factor 1c; NAC019, NAM, ATAF and CUC domain-containing protein 019; ND, neutral day; NO, nitric oxide; npr1-3, nonexpressor of pathogenesis-related genes 1-3; PA, phosphatidic acid; Phe, phenylalanine; PIPK, phosphatidylinositol phosphate kinase; PLD, phospholipase D; Pro, proline; Raf, raffinose; ROS, reactive oxygen species; SA, salicylic acid; *sid2-1*, salicylic acid induction deficient 2-1; WRKY, WRKY domain-containing protein; WT, wild-type.

## 4. Discussion

The metabolic reprogramming caused by the *DAPAT* mutation (Cavalcanti *et al*., 2018; Gouveia *et al*., 2026) and the altered stress responses of *dapat* plants (Neves *et al*., 2024; Gouveia *et al*., 2025, *bioRxiv preprint*) raised the possibility that constitutively elevated free SA levels (Supplementary Fig. S1b), previously linked to enhanced resistance against Pseudomonas syringae (Rate & Greenberg, 2001; Song *et al*., 2004), may also shape heat-stress acclimation. This hypothesis is supported by evidence that SA-dependent signaling contributes to basal thermotolerance in *Arabidopsis* (Clarke *et al*., 2004) and by recent work showing that elevated temperature modulates SA biosynthesis and signaling, thereby linking temperature responses with plant immunity and resilience (Li *et al*., 2025). We therefore used *dapat* as a metabolic– hormonal perturbation to test whether lysine deficiency and SA accumulation interact to reprogram the physiological, metabolic and transcriptional responses to heat stress. Our results provide evidence that *DAPAT* mutation elicits a uniquely distinct heat stress response under both prolonged warming (PW) and heat shock (HS) conditions, revealing genotype-specific patterns in *dapat* that are not only attributable to their SA enrichment (Figure 5). Notably, the recovery-phase data, following PW (Figure 1) and HS (Figure 3), confirm that the lysine biosynthesis impairment does not compromise recovery capacity post-heat exposure.

### DAPAT mutation reveals an alternative route to heat-stress regulation

The analysis of transcript levels in *dapat* under PW and HS conditions (Figure 2) reveals genotype specific response magnitudes. Post-PW induced distinct expression patterns in *HSFA1a, WRKY26*, and *HSP90*, while HS triggered differential responses in *HSFA1b, HSFA1d, bZIP28, DREB2A*, and *NAC055* in mutant plants. These findings reveal distinct transcriptional cascades within the canonical heat-stress response, in which *HSFA1a* acts as a central regulator of the heat-stress response (HSR), coordinating the induction of downstream transcription factors, including *HSFA2, DREB2A* and *MBF1C* (Liu & Charng, 2013; Ohama *et al*., 2016). Notably, the elevated HSP90 expression in *dapat* under control conditions (Figure 2a) suggests a constitutive activation of heat stress machinery, while its attenuation during post-PW recovery (Figure 2c) indicates active resetting mechanisms. In parallel, *dapat* plants showed strong induction of *WRKY26* after PW, with transcript levels increasing 8.3-fold relative to the corresponding control (Figure 2a). Because *WRKY26* is rapidly induced by heat and acts together with *WRKY25* and *WRKY33* to promote thermotolerance, including the expression of heat-shock proteins and ethylene-biosynthesis genes (Li *et al*., 2011), its enhanced expression in *dapat* may reflect an anticipatory response that could contribute to protection during heat exposure.

Under control conditions and following HS induction, *bZIP28* transcript levels were selectively higher in *dapat* plants, suggesting that ER-associated stress signaling may contribute to the mutant response (Gao *et al*., 2008; Liu *et al*., 2007). This interpretation is further supported by the pronounced induction of *HSP101* in *dapat* after HS, consistent with activation of a robust heat-protective program. Alongside pronounced *DREB2A* induction after treatment, which may reflect enhanced transcript or protein stability under heat (Mizoi *et al*., 2019), *NAC055* transcripts also increased in *dapat* (Figure 2b). Given that *NAC055* is co-expressed with *ATAF1* (*ANAC002*) and has been linked to attenuation of thermomemory (Shahnejat Bushehri *et al*., 2012; Alshareef *et al*., 2022), its induction may reflect altered regulation of heat-stress memory in the mutant. To better understand whether the transcriptional effects persisted beyond the acute response of stress, we examined selected stress-responsive genes after a 7-day recovery period, including *MBF1c*, a coactivator previously linked to thermotolerance (Suzuki *et al*., 2008) and stress responses in *Arabidopsis* (Jaimes-Miranda *et al*., 2020). After recovery, the elevated basal *HSP90* expression observed in *dapat* before stress was not maintained, resulting in downregulation of the transcript; whereas *MBF1c* remained higher in *dapat*, reinforcing a sustained and genotype- specific recovery response (Figure 2c).

Based on the differential expression patterns of genes associated with PW and HS responses in the *dapat* mutant, we decided to include the SA-related mutants *sid2-1* (biosynthesis) and *npr1-3* (signaling) to further explore the interplay between lysine biosynthesis, SA accumulation or depletion, and HSR. The expression analysis of HSR genes in WT, *dapat*, *sid2-1*, and *npr1-3* under heat stress (Figure 3) showed pronounced upregulation of *HSFA1a* and *bZIP60* in *dapat* plants under control conditions, suggesting a primed stress-alert state (Cavalcanti *et al*., 2018; Gouveia *et al*., 2025, *bioRxiv preprint*). Remarkably, this priming response was absent in the SA-deficient mutants, particularly *npr1-3*, which, like *dapat*, exhibits elevated basal free SA levels (Supplementary Fig. 4Sa). Notably, upon HS, *dapat* exhibited no differential expression of *HSFA1a*, whereas both *sid2-1* and *npr1-3* demonstrate a robust transcriptional upregulation (Figure 4a). A comparable pattern emerged for *bZIP60*, showing that after HS, the induction was attenuated in *dapat* yet strongly enhanced in the SA mutants (Figure 4a). Activated by *INOSITOL REQUIRING ENZYME1 IRE1*-driven mRNA splicing, *bZIP60* migrates to the nucleus and induces ER chaperones, ER-associated degradation factors, and heat-shock proteins, thereby reinforcing thermotolerance (Nagashima *et al*., 2011; Li *et al*., 2020). Together, these results suggest that SA status shapes both the basal primed state and the acute transcriptional output of the HSR, with *dapat* displaying a distinct regulatory signature compared with the SA-impaired lines.

Furthermore, the downregulation of *DREB2A* in *dapat*, *sid2-1*, and *npr1-3* suggests that this pathway is not the main heat-stress output in these genotypes (Figure 4a). Instead, the induction of *MBF1c* after the HS recovery (Figure 4b), alongside in *dapat* and after PW (Figure 2c), points to an alternative and more sustained transcriptional program that may help define their heat-response signature.

### Salicylic acid–modulated stomatal and electron transport dynamics under heat stress

Following the identification of a distinct transcriptional signature in the *dapat* mutant, gas exchange parameters were quantified in WT, *dapat*, *sid2-1*, and *npr1-3* plants after exposure to PW and HS to determine whether SA status also shaped physiological heat responses. Under control conditions, the SA mutants *sid2-1* and *npr1-3* showed higher stomatal conductance (*gs*) and transpiration (*E*) than WT, whereas *dapat* maintained the lowest basal conductance (Table 1), consistent with a more conservative physiological state (Cavalcanti *et al*., 2018). Upon warming, all genotypes reduced *gs* by approximately 30%, in line with a VPD*leaf*-driven stomatal closure largely independent of SA status and likely dominated by ABA signaling (Zamora *et al*., 2021). This was accompanied by reduced *E*, confirming stomatal limitation of gas exchange under PW (Wang *et al*., 2020). Notably, PSII quantum yield (Φ*PSII*) remained stable across genotypes, indicating that PSII integrity was preserved under moderate heat (Table 1). By contrast, electron transport rate (*ETR*) showed genotype-specific acclimation: WT increased *ETR* modestly, both SA-deficient mutants showed a stronger increase, whereas *dapat* was largely unresponsive, suggesting a trade-off between water conservation and photochemical adjustment (Zhang & Sharkey, 2009).

After HS and recovery, all genotypes showed strong reductions in *gs* an, consistent with a generalized post-stress closure response. In contrast, PSII efficiency remained stable and recovered to WT-like levels in *dapat*, whereas *ETR* was maintained or slightly enhanced in the SA-related mutants (Table 1), suggesting metabolic compensation during recovery. Together, these results indicate that SA status influences not only the transcriptional heat-stress signature, but also the balance between stomatal control and photosynthetic acclimation. Notably, *dapat* adopted a distinctly conservative strategy under heat, prompting us to examine whether this response was associated with altered SA homeostasis.

### Salicylic acid metabolism is dynamically reconfigured during heat stress and recovery

In WT, *dapat*, *sid2-1*, and *npr1-3* plants, PW triggered genotype-specific changes in SA metabolism (Figure 5a). After seven days of warming, all genotypes showed a marked decrease in free SA, consistent with its consumption or turnover during stress and with a role for SA in limiting oxidative damage (Larkindale *et al*., 2000; Tuang *et al*., 2020; Sangwan *et al*., 2022). Strikingly, *dapat* uniquely displayed concomitant reductions in SAG and total SA, together with a compensatory increase in SGE, suggesting that excess SA may be redirected into alternative conjugate pools when homeostasis is challenged (Thompson *et al*., 2017). By contrast, *sid2-1* maintained low SA levels throughout warming and showed little adjustment in SA conjugates, consistent with the role of SID2 (EC 1.14.13.93) for heat-induced SA accumulation. The intermediate phenotype of *npr1-3* supports NPR1 as a signaling component, as reflected by moderate SA depletion and the absence of the strong conjugate shifts seen in *dapat*.

Under HS followed by recovery, *dapat* underwent a rapid depletion of SA, SAG, SEG, and total SA, but these pools largely returned to WT-like levels after recovery (Figure 5b). Notably, *dapat* and *npr1-3* converged on similar levels of SA, SAG, and total SA. However, SEG remained higher in *dapat* than in *npr1-3*, suggesting that NPR1 may contribute to the fine-tuning of SA conjugate balance even after stress release (Liu *et al*., 2020). In summary, these patterns reinforce the idea that heat stress reshapes SA metabolism in a genotype-dependent manner, with *dapat* combining a distinctive transcriptional heat signature, altered gas-exchange behavior, and a marked rewiring of SA pools. These metabolic changes set the stage for the GC-MS analysis of WT, *dapat*, *sid2-1*, and *npr1-3* under PW, HS, and recovery, which aimed to determine how SA- dependent shifts in primary metabolism may contribute to stress acclimation and recovery.

### DAPAT impairment amplifies heat-induced reprogramming of primary metabolism

Primary-metabolism dynamics confirmed genotype-dependent metabolic reprogramming during heat stress. Prior to stress, *dapat* plants display a pre stressed metabolome marked by accumulation of nearly all amino acids analyzed, elevated TCA cycle intermediates, increased sucrose, and depleted reduced-sugars (Cavalcanti *et al*., 2018; Gouveia *et al*., 2026), whereas WT and SA-pathway mutants maintain baseline profiles (Figure 6). During warming, WT undergoes broad depletion of amino acids, consistent with stress induced catabolic shifts. In contrast, *dapat* sustains its pronounced amino acid enrichment (e.g., GABA, lysine, and BCAA) through day 7, whereas *sid2-1* and *npr1-3* shared only lysine accumulation and pronounced proline depletion, consistent with a role for SA signaling in shaping amino-acid homeostasis under heat stress (Liu *et al*., 2010; Khan *et al*., 2015). Across genotypes, warming induces accumulation of organic acids, while 2-OG and pyruvic acid decline markedly, suggesting altered TCA cycle flux under heat stress (Wang *et al*., 2020). Sugar profiles show stable sucrose yet a trend toward hexose depletion in WT and *npr1-3*. Moreover, raffinose remains elevated in *dapat* and *npr1-3*, in line with its protective osmolyte function during heat stress (Nishizawa *et al*., 2008; Egert *et al*., 2013; Reichelt *et al*., 2023).

Metabolic profiling of WT, *dapat*, *sid2-1* and *npr1-3* under HS and 24-h recovery revealed a second layer of genotype-specific shifts (Figure 7) that broadly paralleled those observed under PW. Under control conditions, *dapat* again accumulated multiple amino acids and TCA intermediates, including high citrate/isocitrate levels, while raffinose was depleted. By contrast, *sid2-1* and *npr1-3* showed reduced glutamate, glycine, serine and reduced sugars, together with an elevated citrate/isocitrate ratio, fumarate and malate; raffinose enrichment was observed only in *sid2-1*. HS triggered a global increase in amino acids, particularly β-alanine, BCAA and lysine, alongside a decline in reducing sugars and strong raffinose accumulation in all genotypes. During recovery, BCAA and lysine declined below recovery controls, reducing sugars were restored, and raffinose remained elevated. Notably, HS followed by recovery largely reversed the *dapat*-associated metabolic reprogramming observed under control conditions, restoring amino-acid, sugar and TCA-cycle profiles toward WT-like states (Figure 7).

Together, these data indicate that *dapat* sustains a constitutive, stress-like metabolic state under both control and warming conditions, whereas SA-deficient and SA-signaling mutants display a more restrained but still genotype-specific reprogramming. More broadly, the rapid partial reset of the *dapat* metabolome after HS suggests that this genotype may be metabolically primed for heat exposure yet still retains the capacity to recover during stress relief. Notably, the PCA patterns reinforce that *dapat* undergoes the most extensive heat-induced metabolic reprogramming, linking its primary-metabolism phenotype to altered BCAA and TCA-cycle homeostasis. This response is consistent with the lysine-biosynthesis defect, which drives compensatory BCAA overaccumulation (Cavalcanti *et al*., 2018). By contrast, *sid2-1* and *npr1-3* occupy intermediate positions, indicating that disruptions in SA metabolism also promote metabolic rerouting, but less strongly. Overall, these patterns suggest that the DAPAT mutation amplifies primary-metabolite reprogramming under HS and underscore its role in coordinating BCAA homeostasis at elevated temperature (Bulut *et al*., 2025).

### RNA-seq reveals conserved and genotype-dependent transcriptional responses to heat shock and recovery

Functional annotation of the top variable genes in WT and *dapat* plants exposed to HS and subsequent recovery showed that the transcriptional response extended well beyond canonical HSPs, including epigenetic regulation, amino-acid metabolism, and photosynthesis-related genes, in line with multi-omics evidence that the plant HSR is embedded in a wider network coordinating chromatin state, metabolism, and ROS signaling. The presence of *HSF, DREB*, and *bZIP*-family targets among the variable genes further supports a model in which HSFA1-driven transcription branches into *DREB2A*- and *bZIP28*/*60-*dependent sub-networks that fine-tune specific aspects of the stress response (Ding *et al*., 2020; Haider *et al*., 2022). The transcriptional response was characterized by the coordinated induction of canonical heat-shock proteins and molecular chaperones, including *HSP101, HSP70, HSP83, HSP17.4, HSP17.6, HSP17.8, HSP18.2, HSP23.5/23.6*, and *DNAJ*. This pattern is consistent with the classical model in which *HSFA1* proteins act as master regulators of the HSR, being released from *HSP70/HSP90* repression upon heat exposure and subsequently activating downstream HSFs and HSP genes (Liu *et al*., 2011).

Subsequently, the dataset revealed not only common transcriptomic responses, but also genotype-specific trends. *dapat* exhibited a persistent downregulation of the chloroplastic chlorophyll a-b binding protein *LHCB4*.1 (CP29.1) regardless of the treatment. As a minor antenna protein of photosystem II (PSII), the stable reduction of *LHCB4.1* implies a structural reorganization of the thylakoid grana to limit light harvesting and mitigate potential photo-oxidative damage (de Bianchi *et al*., 2011; Caferri *et al*., 2025), which is highly consistent with the conservative photosynthetic strategy observed in the mutant. This resource-conserving response is further substantiated by the downregulation of *LTP3* (encoding a lipid transfer protein). Specifically, *dapat* plants also showed a highly coordinated, constitutive upregulation of genes central to SA-mediated defense and systemic immunity. This suite of upregulated transcripts includes *CHI, CYP71B15, AtCAPE9, CRK6, AtRLP23, AtBG2, DLO1, AT3G28510, TAT, ARCK1, ATMBP, LCR77*, and *PDF1.3*, alongside a substantial accumulation of *AT1G15125, AT1G27020, AT1G13470, AtHIR2, PR-5, AED1, ATGST1*, and *ATGST11*. The constitutive activation and stress-primed expression of these immunological markers suggest that the metabolic bottleneck imposed by the DAPAT mutation, and the resulting chronic accumulation of free SA, triggers a full systemic defense program even in the absence of biotic pathogen pressure (Gouveia *et al*., 2026; Tian *et al*., 2025). The integration of growth repression, the carbon-nitrogen balance, and defense-related gene expression underscores how lysine biosynthesis acts as a major hormonal-metabolic gatekeeper, wherein endogenous energy constraints crosstalk with hormone signaling to prioritize systemic defense programs at the expense of growth-related processes (Gouveia *et al*., 2026; Tian *et al*., 2025).

This defense-oriented transcriptional configuration is further complemented by *GER3* (encoding germin-like protein 3), the MADS-box transcription factor *AGL68* (MAF5), and the calcium:proton antiporter *CAX3*, which are exclusively upregulated in *dapat* while being downregulated in wild-type. The sustained upregulation of *GER3* points to a specialized role in cell wall remodeling and superoxide dismutase-like reactive oxygen species (ROS) scavenging (Collins *et al*., 2010), whereas the distinct upregulation of *AGL68* and *CAX3* likely indicates epigenetic gating of defense responses and tight cytosolic calcium regulation, respectively, which may serve to maintain homeostasis and buffer the chronic energy stress characteristic of the mutant. Furthermore, the native upregulation of *GASA6* and *GGL22* in *dapat* is consistent with the emerging view that lysine biosynthesis acts as a metabolic checkpoint for gibberellin-mediated growth in *Arabidopsis*, linking perturbations in lysine homeostasis to broader changes in carbon–nitrogen balance and GA-associated transcriptional programs (Gouveia *et al*., 2026). *GASA6* encodes a cell wall-localized integrator of gibberellin, ABA, and glucose signaling pathways to regulate growth and cell expansion (Zhong *et al*., 2015), whereas the guard cell-enriched GDSL lipase *GGL22* plays a key redundant role in stomatal dynamics and plant water relations (Xiao *et al*., 2021). Together, their coordinate abundance suggests that *dapat* recalibrates developmental growth and stomatal behavior in response to disrupted lysine homeostasis, thereby optimizing resource allocation.

Within the HS cluster, *dapat* samples showed visibly stronger induction than WT, in agreement with the RT–qPCR data showed previously (Figure 2 and Figure 4a), with preferential upregulation of *HSFA1*, *HSP*, and folding-related transcripts in the mutant. Specifically in HS, the homeostasis of protein quality control is essential, and was evidenced in *dapat* by the stable expression of the type II phosphoinositide kinase *PI4Kgamma3*, whereas downregulated in WT. Given that *AtPI4Kgamma3* regulates nuclear phosphoinositide signaling, ROS homeostasis, and stress gene expression (Akhter *et al*., 2016), the maintenance of its expression indicates a preserved signaling capacity.

### Distinct recovery-phase reprogramming

In general, recovery samples did not cluster with either control or HS samples, indicating that transcriptional reprogramming persists beyond the acute stress phase rather than reverting immediately to baseline. During recovery, genes linked to SA and defense signaling (*e.g*., *DLO1, ATWRKY70, ACD6, PR-5, PCC1, LURP1* and *AED*) were differentially regulated during heat stress and recovery, supporting the involvement of SA-linked immune programs in the genotype-dependent transcriptional response (Figure 8), and paralleling the known convergence of the HSR with SA, ethylene, and trehalose dependent pathways downstream of *MBF1c*. This transcriptional signature matches with the SA-pool rewiring and amino-acid/TCA-cycle reprogramming already described in *dapat* plants, suggesting that the same genotype tends to couple its metabolic and hormonal adjustments to a SA-linked transcriptional state rather than a rapid return to baseline. The genotype-dependent induction of *MBF1c* during recovery in *dapat* plants is therefore compatible with a role for DAPAT-linked metabolism in modulating this SA-associated branch of thermotolerance (Suzuki *et al*., 2008), rather than the core HSF–HSP chaperone module itself.

Specifically in *dapat*, the stress-responsive marker *KIN1* is downregulated during the recovery phase, reflecting an efficient transcription reset and attenuation of core stress-responsive outputs once the acute thermal pressure is relieved. This post- stress control is also supported by the specific upregulation of the mitochondrial protease *Deg11* in *dapat*, pointing to an enhanced requirement for protein quality control and degradation of damaged proteins in the mitochondrial stroma/lumen during stress release (Schuhmann & Adamska, 2012). Taken together, the transcriptome assessment suggests that the enhanced heat sensitivity of *dapat* reflects not the activation of a distinct pathway but a quantitatively amplified engagement of the same core *HSFA1*–*HSP* module that governs the *Arabidopsis* HSR, with additional divergence emerging in the SA/*MBF1c*-associated recovery response, a transcriptional signature that parallels the distinctive gas-exchange, SA, and primary-metabolism responses (Figure 9) of *dapat* compared with WT and the SA-pathway mutants.

## 5. Conclusion

Taken together, our results show that DAPAT dysfunction reshapes heat-stress acclimation through an integrated reprogramming of primary metabolism, salicylic acid homeostasis, and transcriptional control. Rather than activating a novel heat-response pathway, *dapat* displays a primed engagement of the conserved *HSFA1*–*HSP* module, accompanied by a distinct recovery-phase signature that involves SA- and *MBF1c*- associated regulation. This pattern is consistent with the notion that impaired lysine biosynthesis places the mutant in a preconditioned, stress-alert state that alters how heat signals are perceived, amplified, and resolved. At the physiological level, *dapat* plants combine conservative gas-exchange behavior with preserved photochemical performance and an unusual SA profile, reinforcing the idea that DAPAT mutation affects the balance between protection and growth. At the metabolic level, the mutant sustains strong accumulation of amino acids, TCA-cycle intermediates and compatible solutes under heat, while still retaining the capacity to partially reset its metabolome during recovery.

These features indicate that DAPAT deficiency does not abolish recovery competence but instead redirects carbon and nitrogen partitioning toward a distinct heat- adaptive state. The transcriptome data further reveals that this reprogramming extends beyond canonical heat-shock genes to include developmental, defense, and stomatal regulators such as *GASA6* and *GGL22*, whose basal upregulation is consistent with a broader role for lysine biosynthesis as a metabolic checkpoint for gibberellin-mediated growth and resource allocation. In this context, the DAPAT mutation appears to couple metabolic limitation to hormonal and developmental remodeling, while also maintaining a heightened defense-like signature. Altogether, our study identifies lysine biosynthesis as a point of integration linking metabolism, SA signaling, growth regulation, and thermotolerance in *Arabidopsis*.

## Acknowledgments

This research was funded by the National Council for Scientific and Technological Development (CNPq Brazil, Grant 407276/2021 1), the National Institute of Science and Technology in Plant Stress Physiology (INCT Plant Stress Physiology/CNPq Brazil, Grant 406455/2022 8), and the FAPEMIG (Foundation for Research Assistance of the Minas Gerais State, Brazil, Grant RED 00053 16). Scholarships and research fellowships from CNPq Brazil to WLA and ANN are also acknowledged.

## Supplementary Figure Legends

**Supplementary Figure S1. Disruption of chloroplastic lysine biosynthesis, salicylic acid accumulation and experimental design for heat-stress profiling.** (a) Schematic representation of the chloroplastic aspartate-derived lysine-biosynthesis pathway. The pathway shows the conversion of tetrahydrodipicolinate (THDP) to *L,L*-diaminopimelate (*L,L*-DAP), catalysed by *L,L*-diaminopimelate aminotransferase (DAPAT), and the feedback inhibition of dihydrodipicolinate synthase (DHDPS) by *L*-lysine (red dashed line). Reduced DAPAT activity in the *dapat* mutant disrupts lysine biosynthesis and is associated with metabolic alterations, dwarfism and altered leaf morphology relative to wild-type (WT/Col-0) plants. (b) Total salicylic acid (SA) abundance in WT and *dapat* plants (n = 4; two-tailed Student’s *t*-test). The diagram illustrates the proposed model in which SA accumulation in *dapat* may influence the heat-stress response by modulating nuclear transcription factors, stress-responsive genes, antioxidant defenses and reactive oxygen species (ROS) homeostasis. The centre line indicates the mean, box limits indicate the 25th and 75th percentiles, whiskers indicate the minimum and maximum values, and individual points represent biological replicates. (c) Schematic overview of the experimental workflow using four *Arabidopsis* genotypes: WT, *dapat*, *npr1-3* and *sid2-1*. Seedlings were grown on plates for 14 days before transplantation to soil (0 days after transplanting, DAT) and were subsequently subjected to prolonged warming, heat shock (38 °C for 6 h) and post-stress recovery. Experiments were conducted under short-day conditions (12 h light at 20 °C/12 h dark at 20 °C) or neutral-day conditions (12 h light at 22 °C/12 h dark at 20 °C). Tissue-collection time points are indicated by scissors. **Abbreviations:** WT, wild-type; *dapat*, *dapat* mutant line; *npr1-3*, *nonexpressor of pathogenesis-related genes 1-3*; *sid2-1*, *salicylic acid induction deficient 2-1*; AK, aspartate kinase; ASDH, aspartate-semialdehyde dehydrogenase; DHDPS, dihydrodipicolinate synthase; DHDPR, dihydrodipicolinate reductase; DAPAT, diaminopimelate aminotransferase; DAPE, diaminopimelate epimerase; DAPDC, diaminopimelate decarboxylase; *L*-Asp, *L*-aspartate; *L*-ASA, *L*-aspartate semialdehyde; HTPA, 4-hydroxy-2,3,4,5-tetrahydrodipicolinate; THDP, 2,3,4,5- tetrahydrodipicolinate; *L,L*-DAP, *L,L*-diaminopimelate; *meso*-DAP, *meso*- diaminopimelate; SA, salicylic acid; FW, fresh weight; DAT, days after transplanting; ROS, reactive oxygen species; HSR, heat-stress response.

**Supplementary Figure S2. Maximum quantum yield of photosystem II remains stable during prolonged warming and recovery.** (a,b) Maximum quantum yield of photosystem II (*Fv/Fm*) measured in leaves of 6-week-old *Arabidopsis* wild-type (WT), *dapat*, *npr1-3* and *sid2-1* plants under control conditions and after 7 days of prolonged warming (a), followed by a 7-day recovery period (b). Bars represent means ± s.e.m. of independent biological replicates (n = 7). Different lowercase letters indicate significant differences among genotypes and treatments, as determined by two-way ANOVA followed by Tukey’s multiple-comparisons test (P < 0.05). (c) Relative transcript abundance of *DAPAT gene (also named AGD2*, At4g33680) in WT plants under control and warming conditions (P = 0.001, two-tailed Student’s *t*-test). (d) Relative *DAPAT* transcript abundance in WT and *dapat* plants under control and heat- shock conditions. For c and d, centre lines indicate the means, box limits indicate the 10th–90th percentile range, and individual points represent biological replicates (n = 3). Different lowercase letters in d indicate significant differences among genotypes and treatments, as determined by two-way ANOVA followed by Tukey’s multiple- comparisons test (P < 0.05). **Abbreviations:** *Fv/Fm*, maximum quantum yield of photosystem II; WT, wild-type; *dapat*, *dapat* mutant line; *npr1-3*, *nonexpressor of pathogenesis-related genes 1-3*; *sid2-1*, *salicylic acid induction deficient 2-1*; AGD2, aberrant growth and development 2.

**Supplementary Figure S3. Temporal accumulation of salicylic acid and its conjugated derivatives during prolonged warming.** (a–d) Absolute abundance of free salicylic acid (SA) (a), salicylic acid glucose ester (SGE) (b), salicylic acid 2-O-β-D- glucoside (SAG) (c) and the total SA pool (d) in wild-type (WT), *dapat*, *npr1-3* and *sid2-1* plants under control and warming conditions over a 7-day time course, with measurements collected on days 0, 2, 4 and 7. Bars represent means ± s.e.m. of independent biological replicates (n = 4). Different lowercase letters indicate significant differences among genotypes and treatments within each time point, as determined by two-way ANOVA followed by Tukey’s multiple-comparisons test (P < 0.05). **Abbreviations:** WT, wild-type; *dapat*, *dapat* mutant line; *npr1-3*, *nonexpressor of pathogenesis-related genes 1-3*; *sid2-1*, *salicylic acid induction deficient 2-1*; SA, free salicylic acid; SGE, salicylic acid glucose ester; SAG, salicylic acid 2-O-β-D-glucoside; total SA, total salicylic acid pool; FW, fresh weight; s.e.m., standard error of the mean.

**Supplementary Figure S4. Accumulation of salicylic acid and its conjugated derivatives during heat shock and recovery.** (a–d) Absolute abundance of free salicylic acid (SA) (a), salicylic acid glucose ester (SGE) (b), salicylic acid 2-O-β-D-glucoside (SAG) (c) and the total SA pool (d) in wild-type (WT), *dapat*, *npr1-3* and *sid2-1* plants exposed to heat shock (HS) and subsequent recovery. For each metabolite, comparisons are shown between control and HS treatments (left subpanels) and between control recovery and post-HS recovery (right subpanels). Bars represent means ± s.e.m. of independent biological replicates (n = 4). Different lowercase letters indicate significant differences among genotypes and treatments within each experimental set, as determined by two-way ANOVA followed by Tukey’s multiple-comparisons test (P < 0.05). **Abbreviations:** WT, wild-type; *dapat*, *dapat* mutant line; *npr1-3*, *nonexpressor of pathogenesis-related genes 1-3*; *sid2-1*, *salicylic acid induction deficient 2-1*; SA, free salicylic acid; SGE, salicylic acid glucose ester; SAG, salicylic acid 2-O-β-D-glucoside; total SA, total salicylic acid pool; FW, fresh weight; s.e.m., standard error of the mean.

**Supplementary Figure S5. Principal component analysis of metabolite profiles in wild-type and mutant plants during prolonged warming, heat shock and recovery.** (a) PCA score plot of metabolite profiles during prolonged warming over a 7-day time course. PC1 and PC2 explain 33.5% and 19.3% of the total variance, respectively. Colours denote genotypes, point shapes indicate sampling days (0, 2, 4 and 7), and point sizes distinguish control and warming treatments. (b) PCA score plot of primary-metabolite profiles under heat-shock and recovery conditions. PC1 and PC2 explain 34.4% and 27.1% of the total variance, respectively. Colours denote genotypes, and point shapes indicate control, control-recovery, heat-shock and recovery conditions. **Abbreviations:** PCA, principal-component analysis; PC, principal component; WT, wild type; *dapat*, *dapat* mutant line; *npr1-3*, *nonexpressor of pathogenesis-related genes 1-3*; *sid2-1*, *salicylic acid induction deficient 2-1*.

**Supplementary Figure S6. Quality assessment, normalization comparison and hierarchical clustering of RNA-seq samples.** (a) Violin and box plots showing gene-expression distributions across individual biological replicates (n = 3) of wild-type (WT) and *dapat* plants under control (Control.Heat), heat-shock (Heat.Shock), control-recovery (Control.Recovery) and post-heat-shock-recovery (Heat.Recovery) conditions. Distributions are shown for the raw counts and after regularized log transformation (rlog) or variance-stabilizing transformation (VST). Centre lines within the violins indicate the medians, and box limits indicate the interquartile ranges. (b) Sample-to-sample distance matrix and hierarchical clustering based on Euclidean distances calculated from VST-transformed read counts. Colour intensity represents distance values ranging from 0 (dark red, greatest similarity) to 100 (light peach, greatest dissimilarity). The annotation bars indicate treatment and genotype. **Abbreviations:** WT, wild-type; *dapat*, *dapat* mutant line; RNA-seq, RNA sequencing; rlog, regularized logarithm transformation; VST, variance-stabilizing transformation.

**Supplementary Figure S7. Principal component analysis of global transcriptomic profiles during heat shock and recovery.** PCA score plot of variance-stabilizing-transformed (VST) RNA-seq expression data based on the 1,000 most variable genes in wild-type (WT) and *dapat* plants. PC1 and PC2 explain 60% and 15% of the total variance, respectively. Point shapes denote genotypes (WT, circles; *dapat*, triangles), and fill colours indicate treatment conditions: control (Control.Heat, grey), heat shock (Heat.Shock, pink), control recovery (Control.Recovery, light blue) and post-heat-shock recovery (Heat.Recovery, light yellow). Individual points represent biological replicates (n = 3). **Abbreviations:** PCA, principal-component analysis; PC, principal component; VST, variance-stabilizing transformation; WT, wild-type; *dapat*, *dapat* mutant line; RNA-seq, RNA sequencing.

**Supplementary Figure S8. Functional and gene-set enrichment analyses of heat- shock-induced transcriptional responses in wild-type and *dapat* plants.** (a–c) Functional profiling of differentially expressed genes in wild-type (WT) plants exposed to heat shock relative to control conditions (WT_Heat.Shock_vs_Control.Heat). Dot plots show enriched Gene Ontology Biological Process (GO BP) terms (a) and KEGG pathways (b). Dot size indicates the number of genes assigned to each term, whereas colour represents the false discovery rate (FDR)-adjusted *P* value. The GeneRatio axis denotes the proportion of input genes associated with each term. (c) Density plots from gene-set enrichment analysis (GSEA) showing enrichment-score distributions for the leading GO BP terms in WT plants. (d–f) Corresponding analyses for *dapat* plants exposed to heat shock relative to control conditions (*dapat*_Heat.Shock_vs_Control.Heat), showing enriched GO BP terms (d), KEGG pathways (e) and GSEA enrichment-score distributions for the leading GO BP categories (f). **Abbreviations:** WT, wild-type; *dapat*, *dapat* mutant line; GO, Gene Ontology; BP, biological process; KEGG, Kyoto Encyclopedia of Genes and Genomes; GSEA, gene-set enrichment analysis; FDR, false discovery rate.

**Supplementary Figure S9. Functional and gene-set enrichment analyses of post- heat-shock recovery responses in wild-type and *dapat* plants.** (a–c) Functional profiling of differentially expressed genes in wild-type (WT) plants during post-heat- shock recovery relative to control recovery conditions (WT_Heat.Recovery_*vs*_Control.Recovery). Dot plots show enriched Gene Ontology Biological Process (GO BP) terms (a) and KEGG pathways (b). Dot size indicates the number of genes assigned to each term, whereas colour represents the false discovery rate (FDR)-adjusted *P* value. The GeneRatio axis denotes the proportion of input genes associated with each term. (c) Density plots from gene-set enrichment analysis (GSEA) showing enrichment-score distributions for the leading GO BP terms in WT plants. (d– f) Corresponding analyses for *dapat* plants during post-heat-shock recovery relative to control recovery conditions (*dapat*_Heat.Recovery_*vs*_Control.Recovery), showing enriched GO BP terms (d), KEGG pathways (e) and GSEA enrichment-score distributions for the leading GO BP categories (f). **Abbreviations:** WT, wild-type; *dapat*, *dapat* mutant line; GO, Gene Ontology; BP, biological process; KEGG, Kyoto Encyclopedia of Genes and Genomes; GSEA, gene-set enrichment analysis; FDR, false discovery rate.

**Supplementary Figure S10. Functional and gene-set enrichment analyses of basal and recovery transcriptomic differences between *dapat* and wild-type plants.** (a–c) Functional profiling of differentially expressed genes in *dapat* relative to wild-type (WT) plants under control conditions (*dapat*_Control.Heat_*vs*_WT_Control.Heat). Dot plots show enriched Gene Ontology Biological Process (GO BP) terms (a) and KEGG pathways (b). Dot size indicates the number of genes assigned to each term, whereas colour represents the false discovery rate (FDR)-adjusted *P* value. The GeneRatio axis denotes the proportion of input genes associated with each term. (c) Density plots from gene-set enrichment analysis (GSEA) showing enrichment-score distributions for the leading GO BP terms. (d–f) Corresponding functional enrichment analyses for *dapat* relative to WT under control-recovery conditions (*dapat*_Control.Recovery_*vs*_WT_Control.Recovery), showing enriched GO BP terms (d), KEGG pathways (e) and GSEA enrichment-score distributions for the leading GO BP categories (f). **Abbreviations:** WT, wild-type; *dapat*, *dapat* mutant line; GO, Gene Ontology; BP, biological process; KEGG, Kyoto Encyclopedia of Genes and Genomes; GSEA, gene-set enrichment analysis; FDR, false discovery rate.

**Supplementary Figure S11. Functional and gene-set enrichment analyses of heat- shock and post-stress-recovery transcriptomic differences between *dapat* and wild- type plants.** (a–c) Functional profiling of differentially expressed genes in *dapat* relative to wild-type (WT) plants under heat-shock conditions (*dapat*_Heat.Shock_*vs*_WT_Heat.Shock). Dot plots show enriched Gene Ontology Biological Process (GO BP) terms (a) and KEGG pathways (b). Dot size indicates the number of genes assigned to each term, whereas colour represents the false discovery rate (FDR)-adjusted *P* value. The GeneRatio axis denotes the proportion of input genes associated with each term. (c) Density plots from gene-set enrichment analysis (GSEA) showing enrichment-score distributions for the leading GO BP terms. (d–f) Corresponding analyses for *dapat* relative to WT plants during post-heat-shock recovery (*dapat*_Heat.Recovery_*vs*_WT_Heat.Recovery), showing enriched GO BP terms (d), KEGG pathways (e) and GSEA enrichment-score distributions for the leading GO BP categories (f). **Abbreviations:** WT, wild-type; *dapat*, *dapat* mutant line; GO, Gene Ontology; BP, biological process; KEGG, Kyoto Encyclopedia of Genes and Genomes; GSEA, gene-set enrichment analysis; FDR, false discovery rate.

