## Supplementary figures and images for "Lysine biosynthesis impairment shapes heat-stress acclimation through metabolic and transcriptional reprogramming in *Arabidopsis thaliana*"

### Figure S1

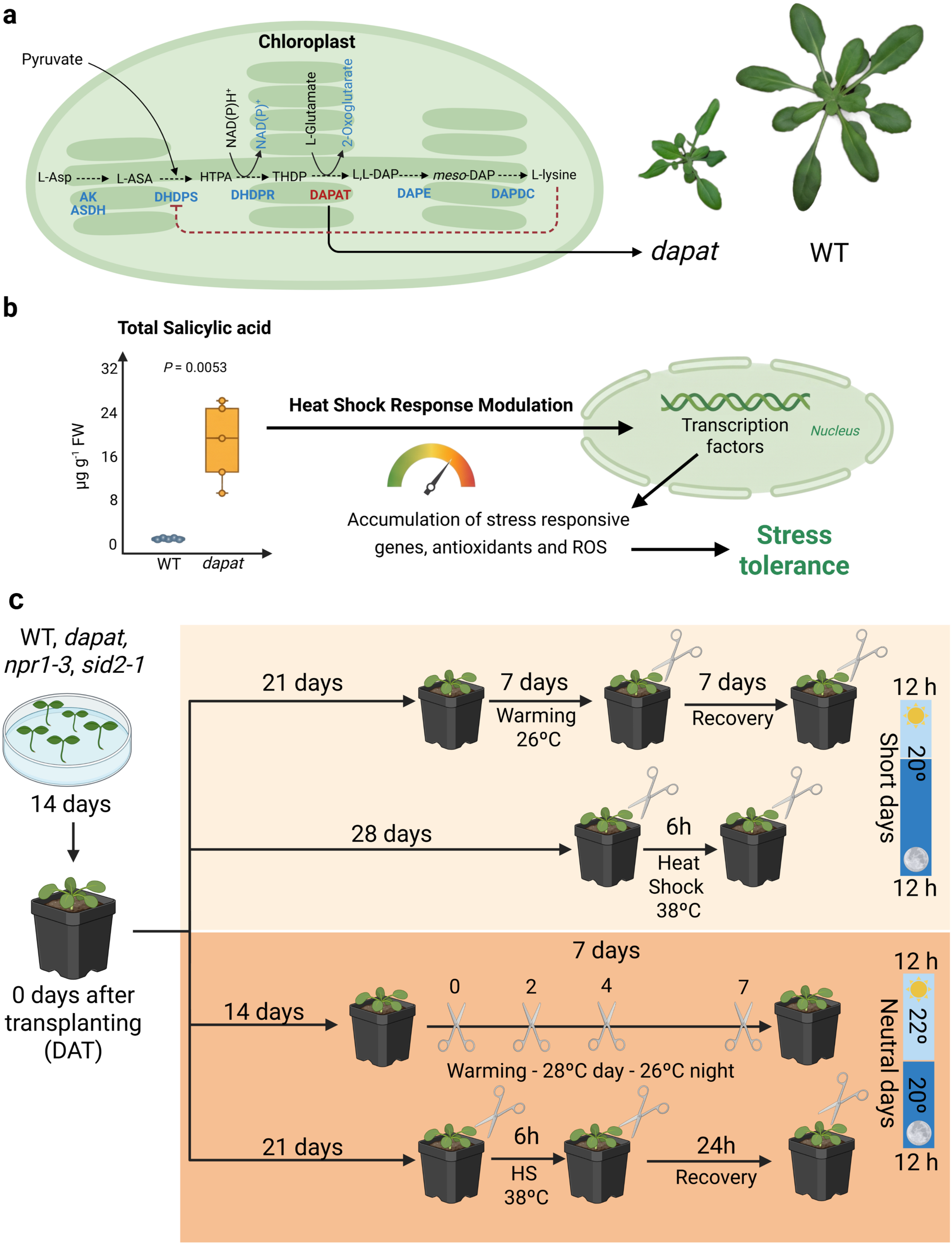

### Figure S2

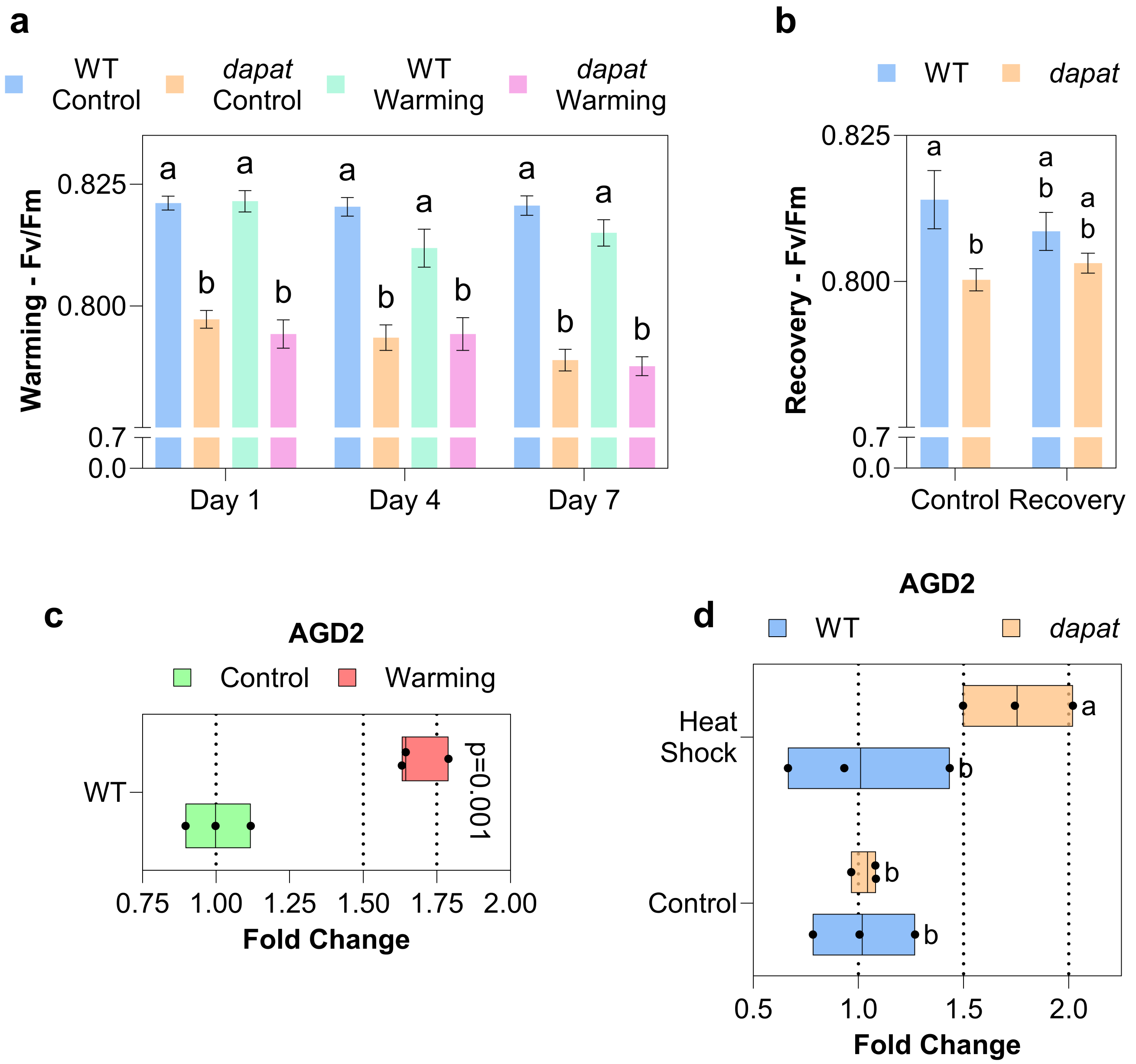

### Figure S3

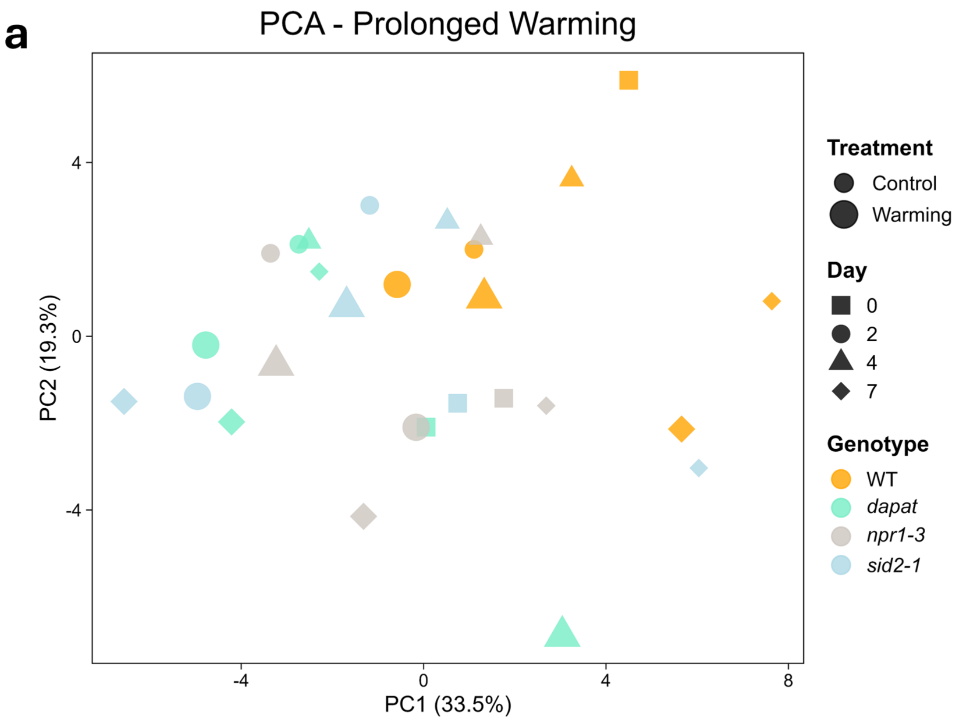

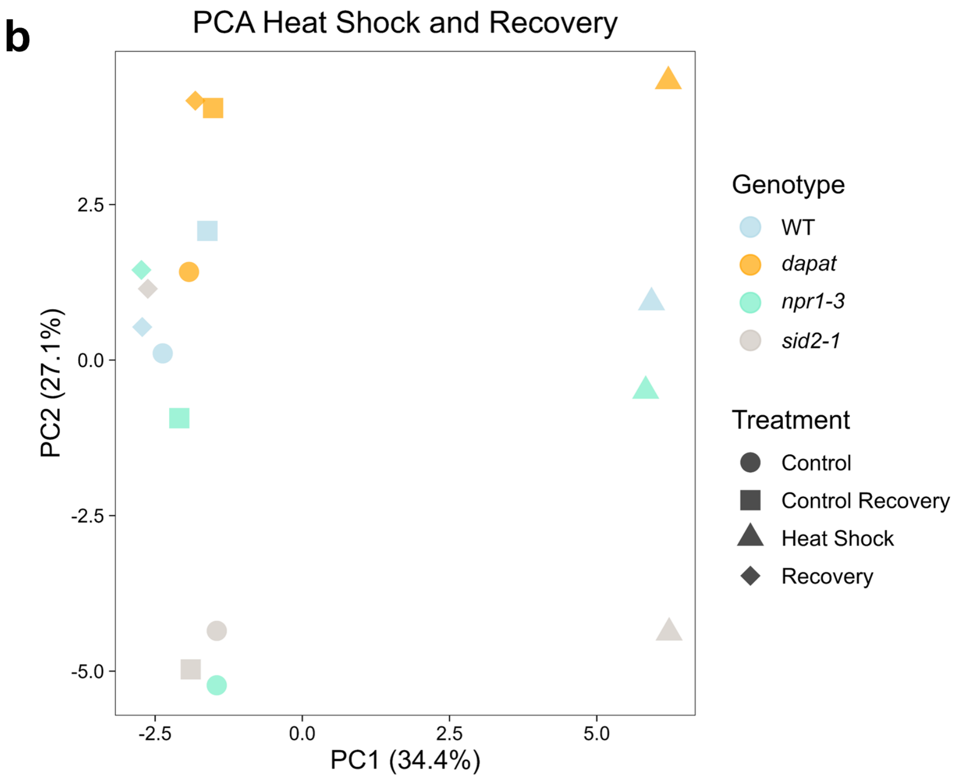

### Figure S4

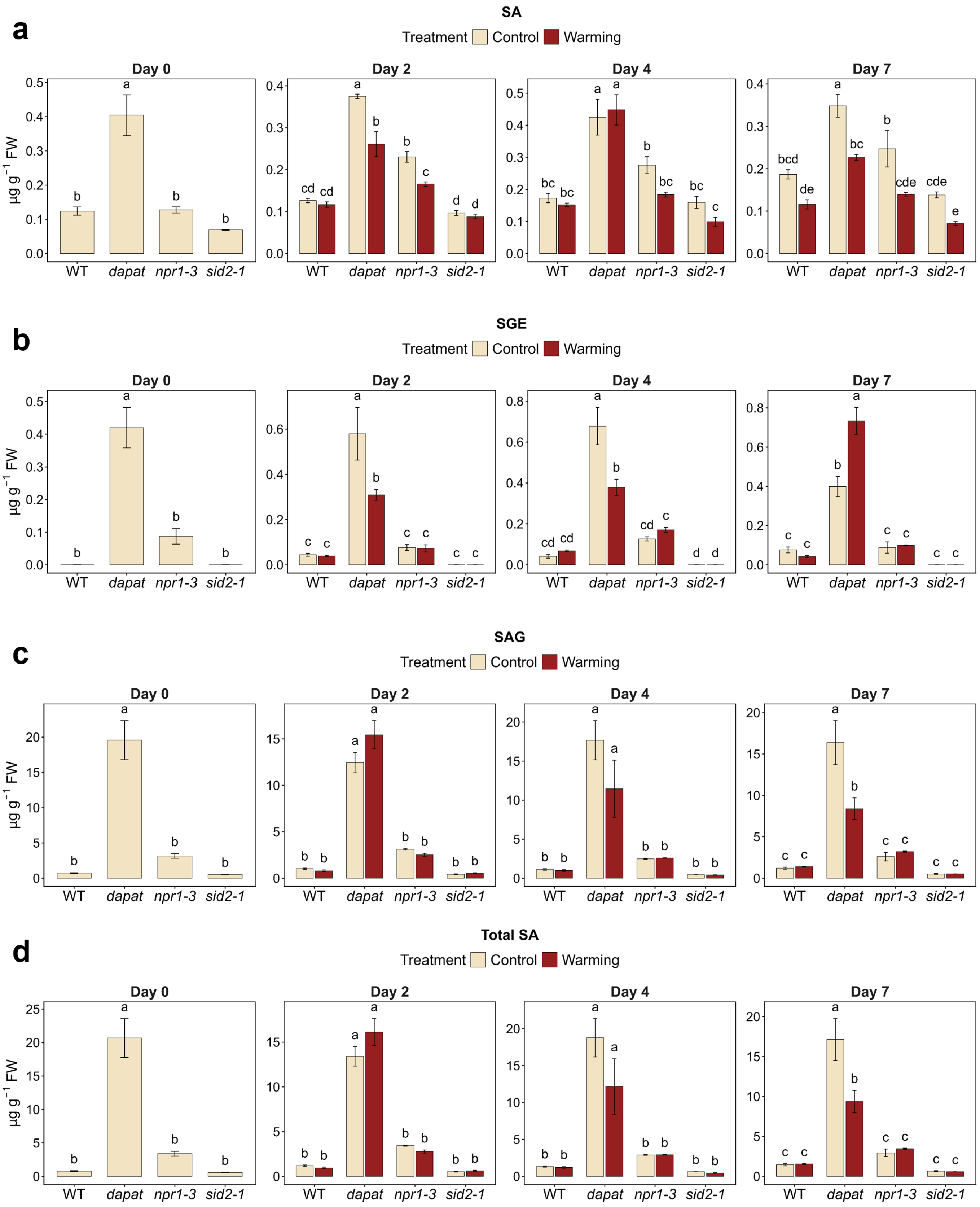

### Figure S5

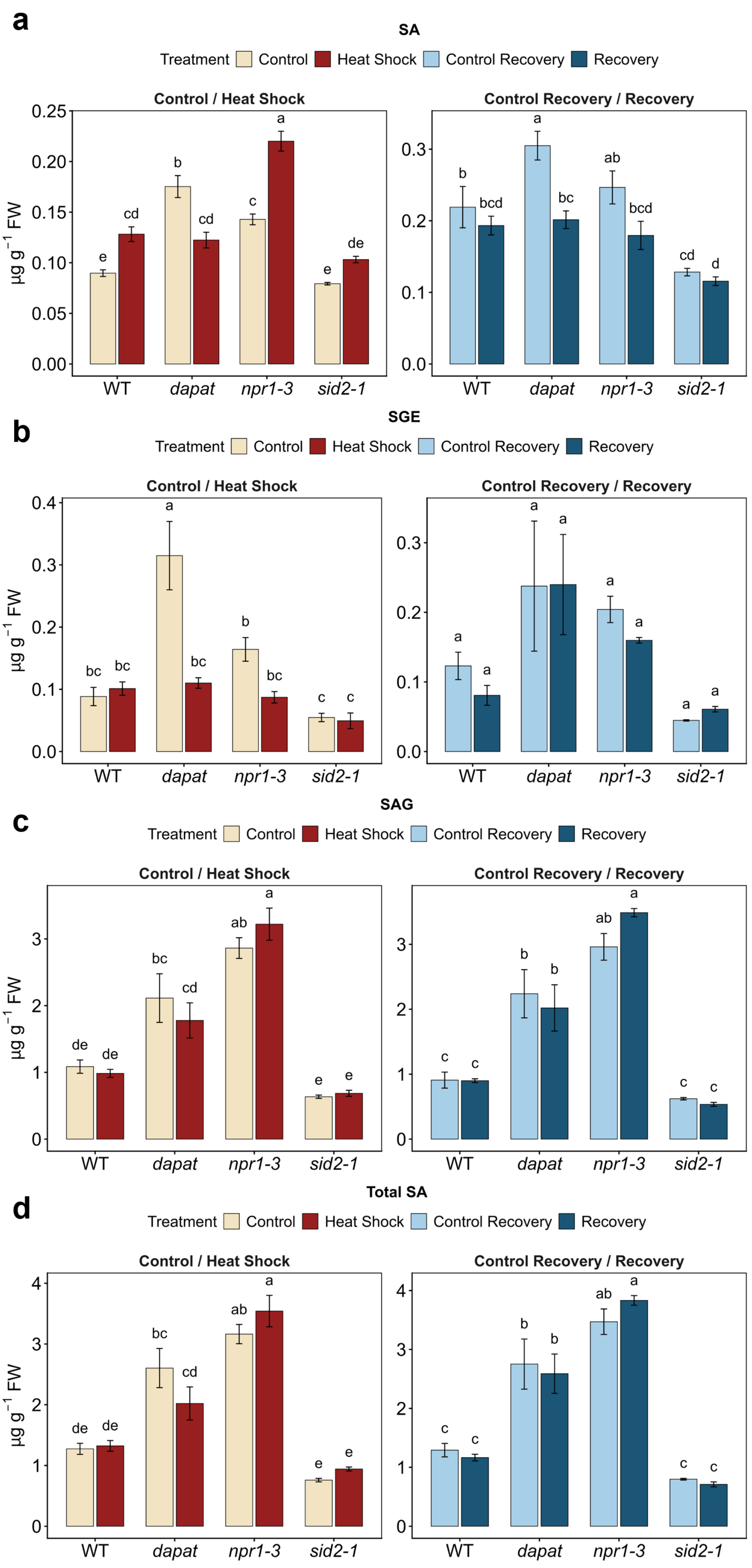

### Figure S6

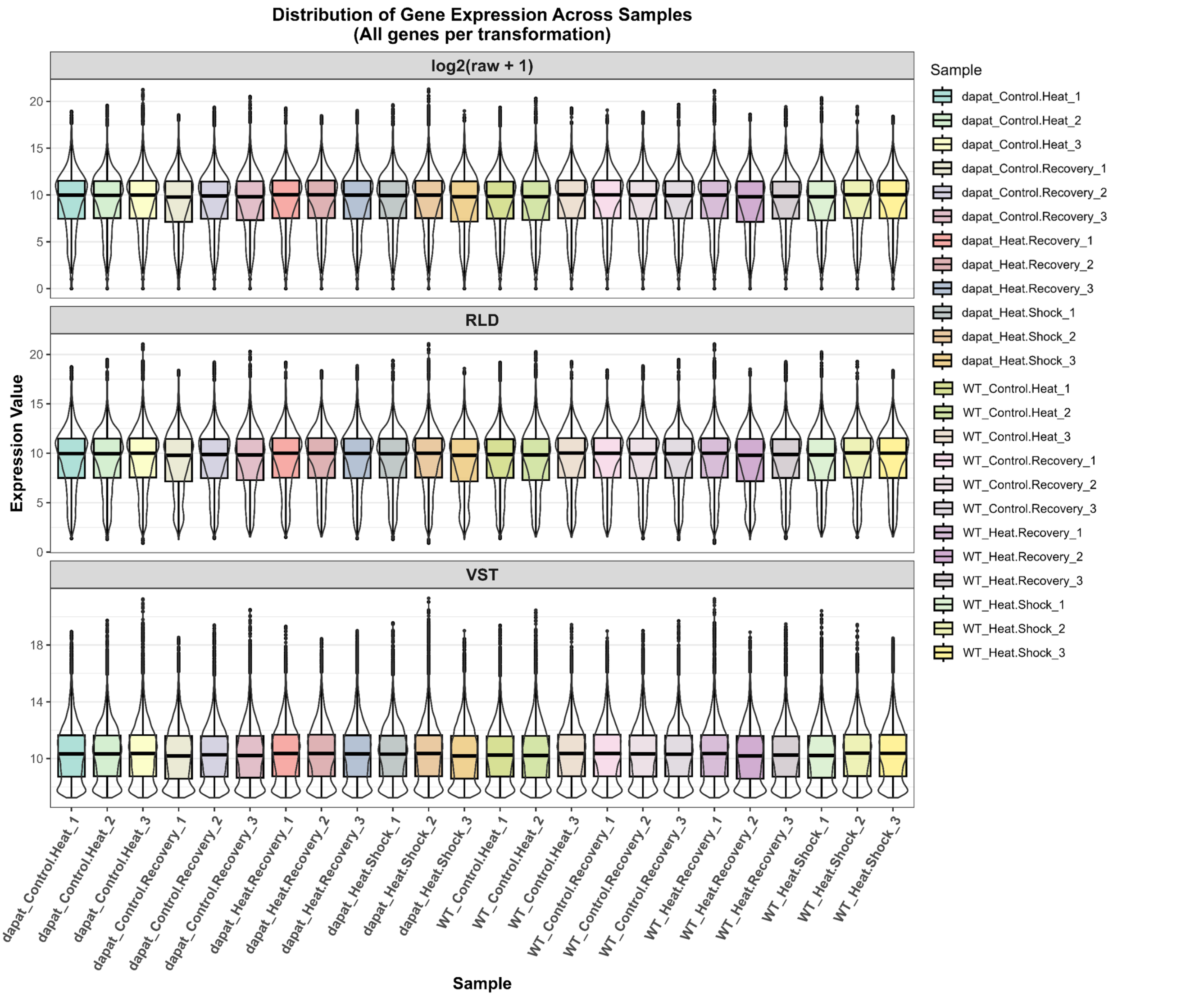

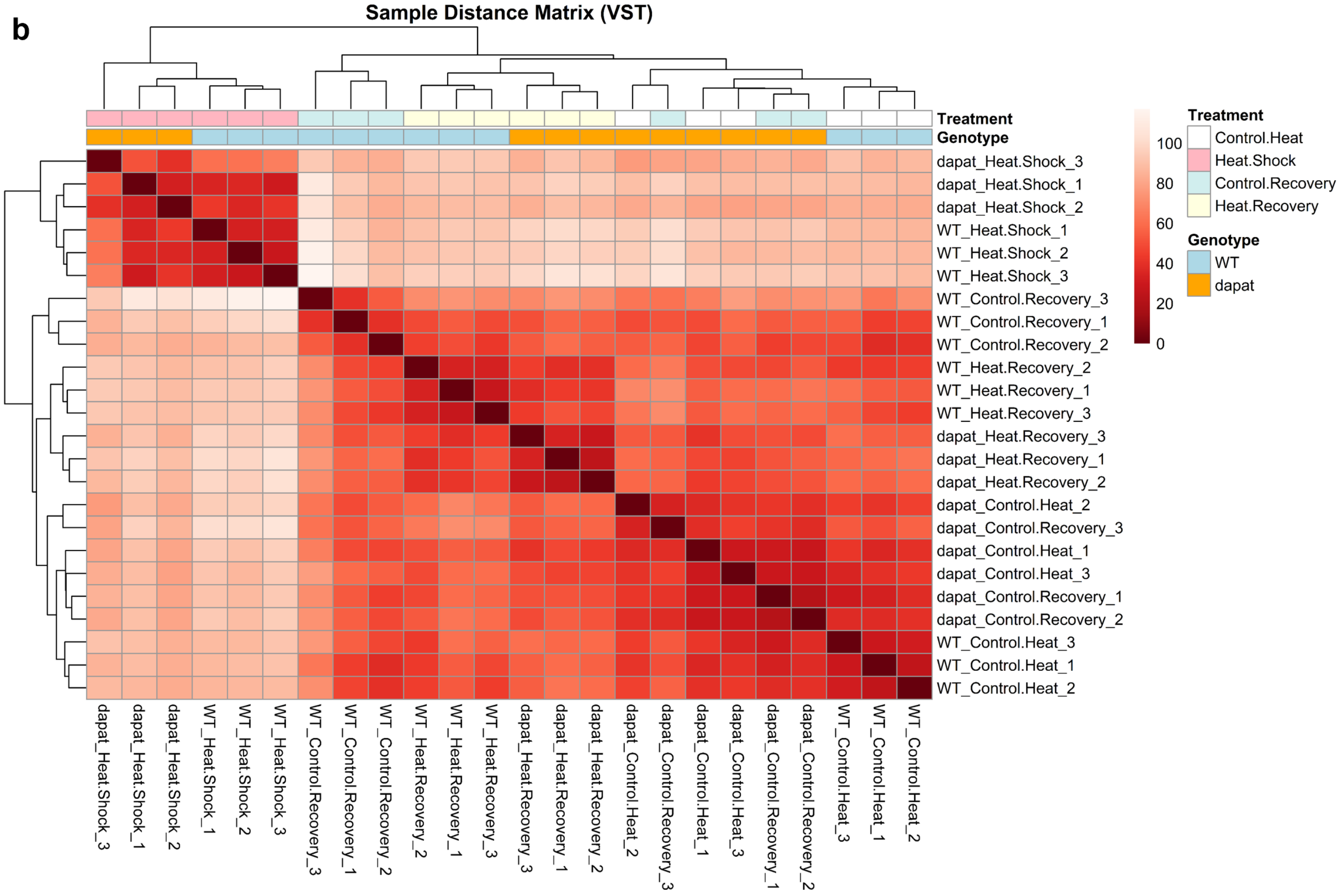

### Figure S7

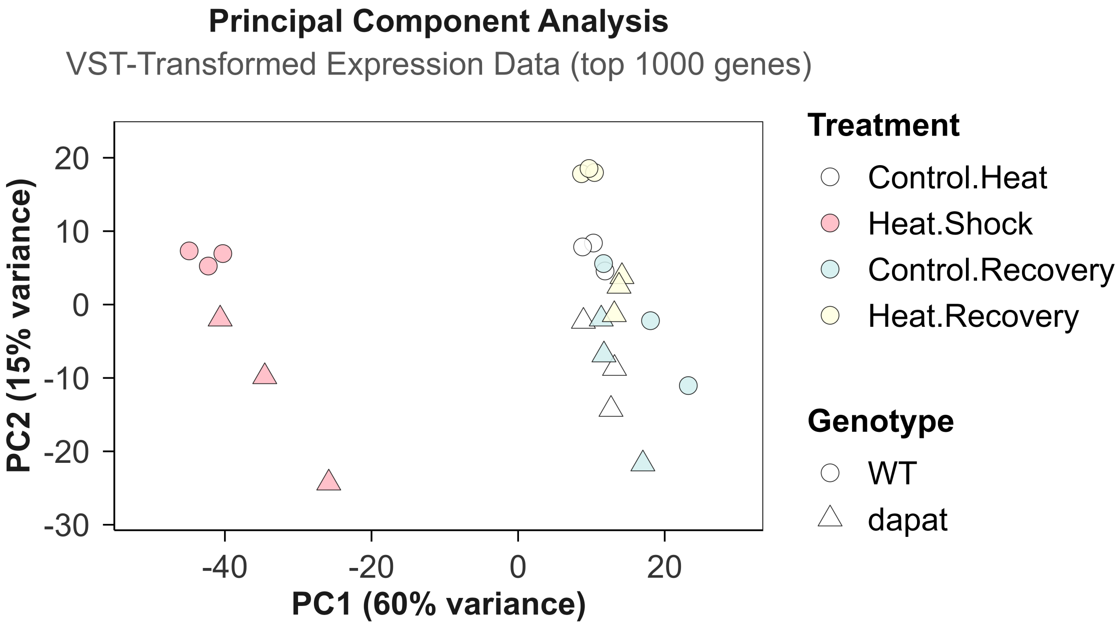

### Figure S8

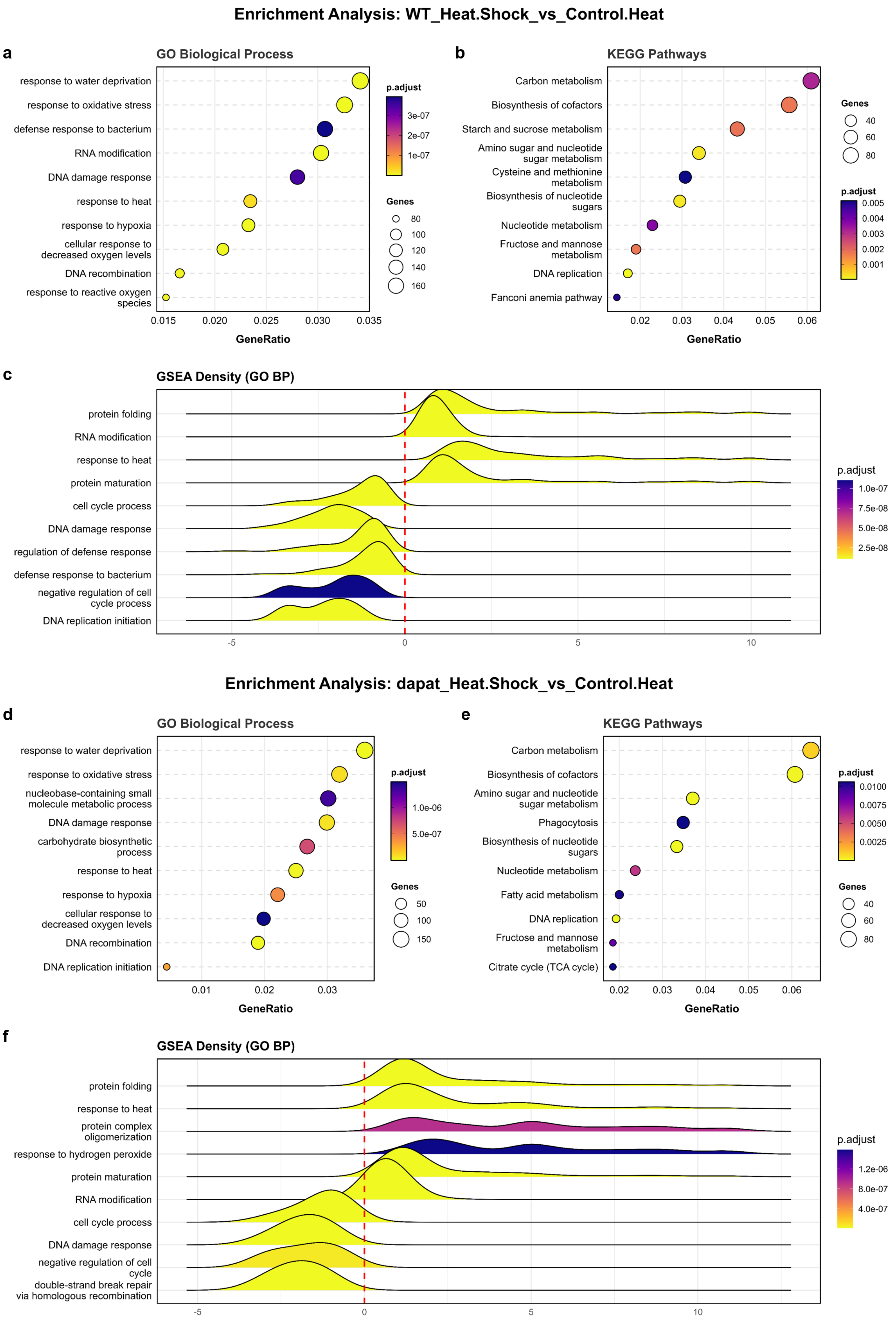

### Figure S10

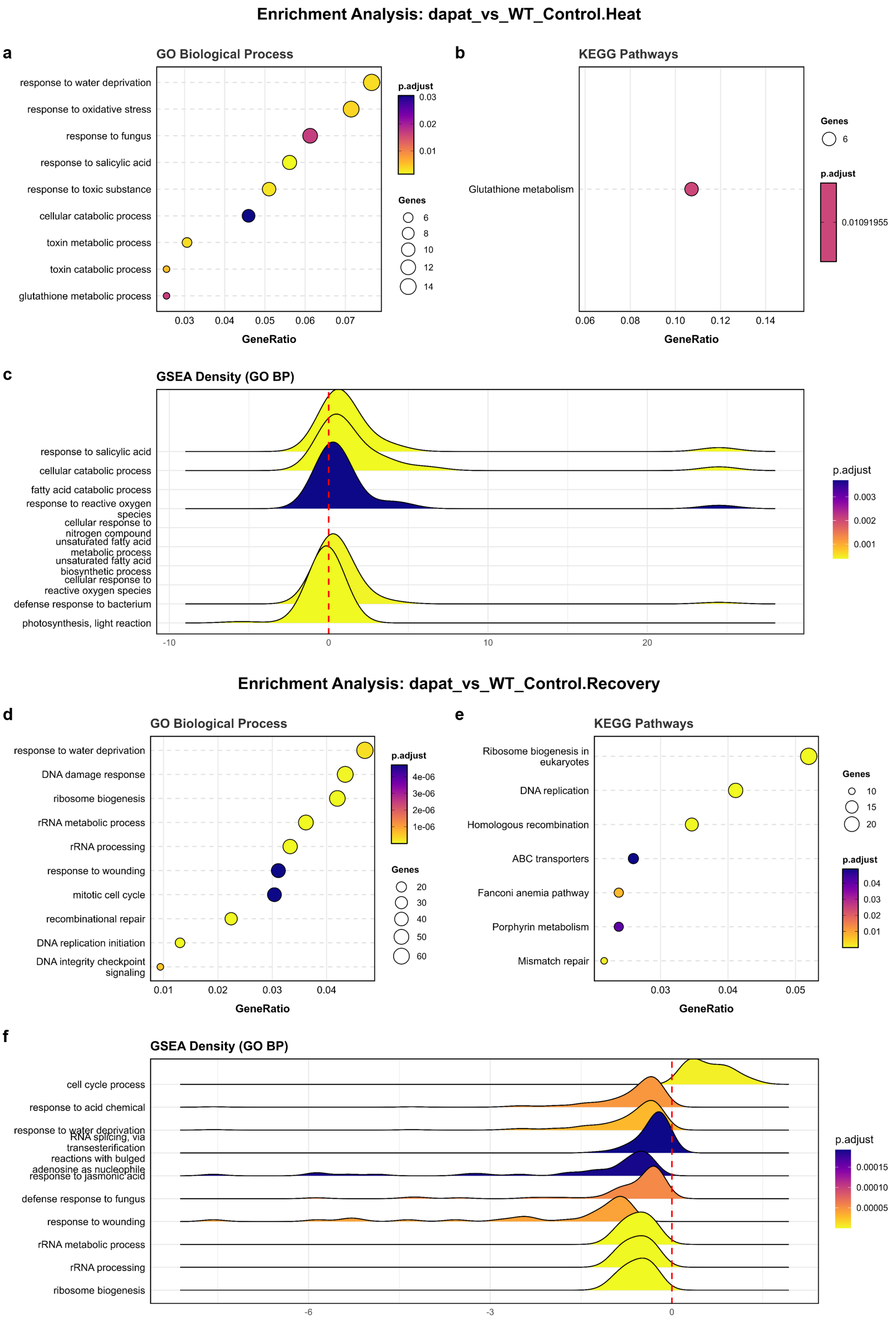

### Figure S11

#### Enrichment Analysis: dapat_vs_WT_Heat.Shock

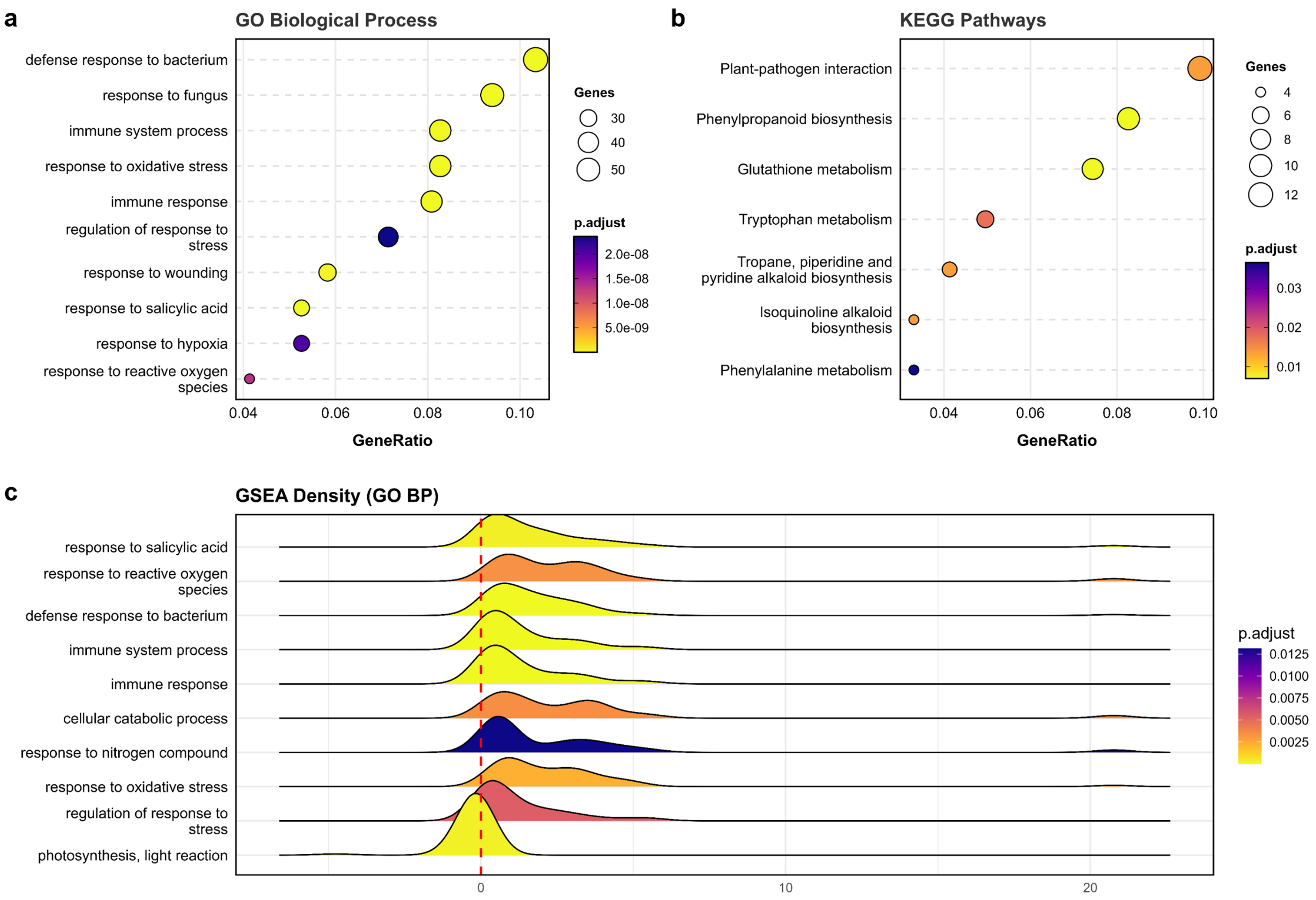


#### Enrichment Analysis: dapat_vs_WT_Heat.Recovery

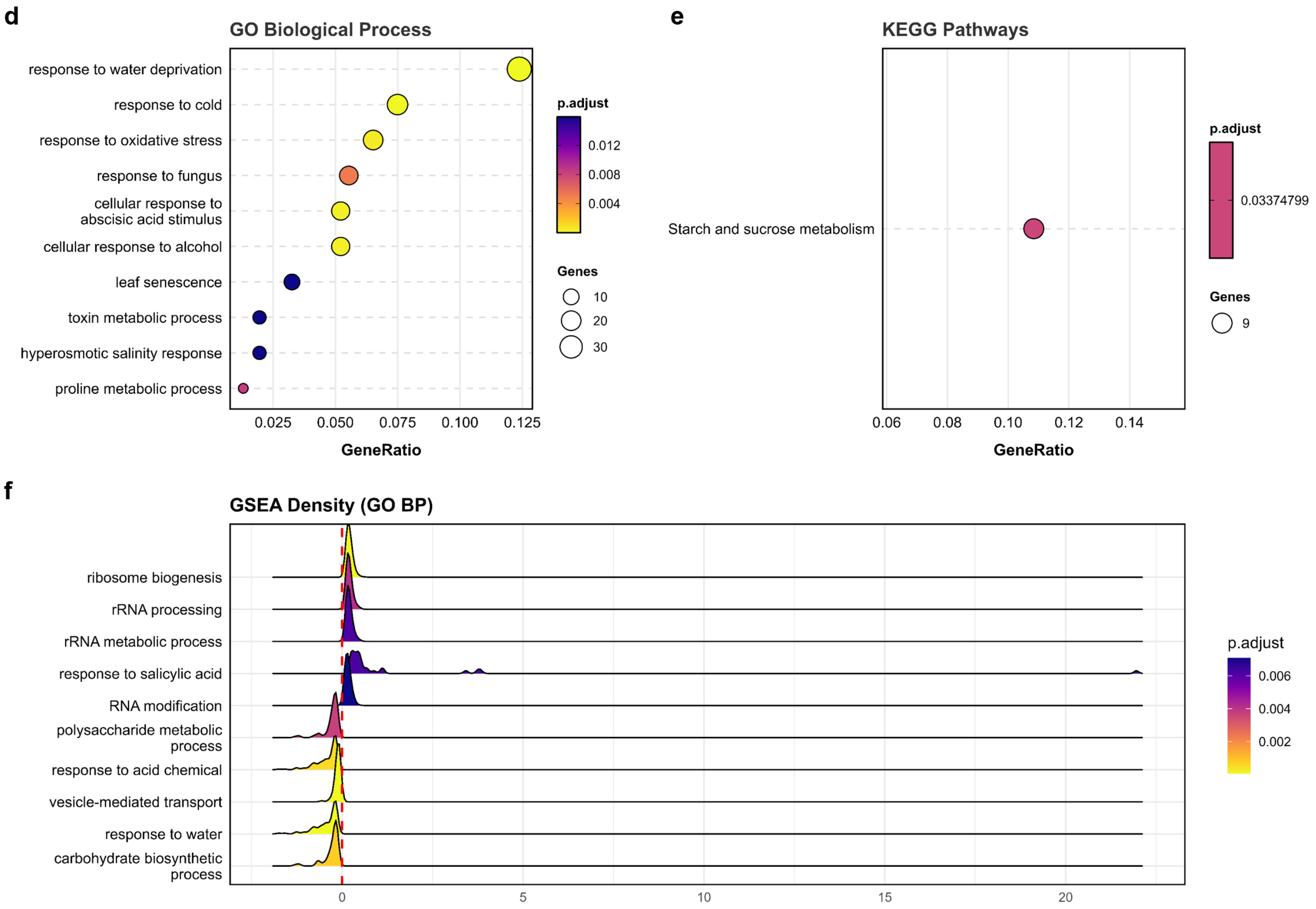
