## Supplementary material for "Lysine biosynthesis impairment shapes heat-stress acclimation through metabolic and transcriptional reprogramming in *Arabidopsis thaliana*": Figure S9

### Enrichment Analysis: WT_Heat.Recovery_vs_Control.Recovery

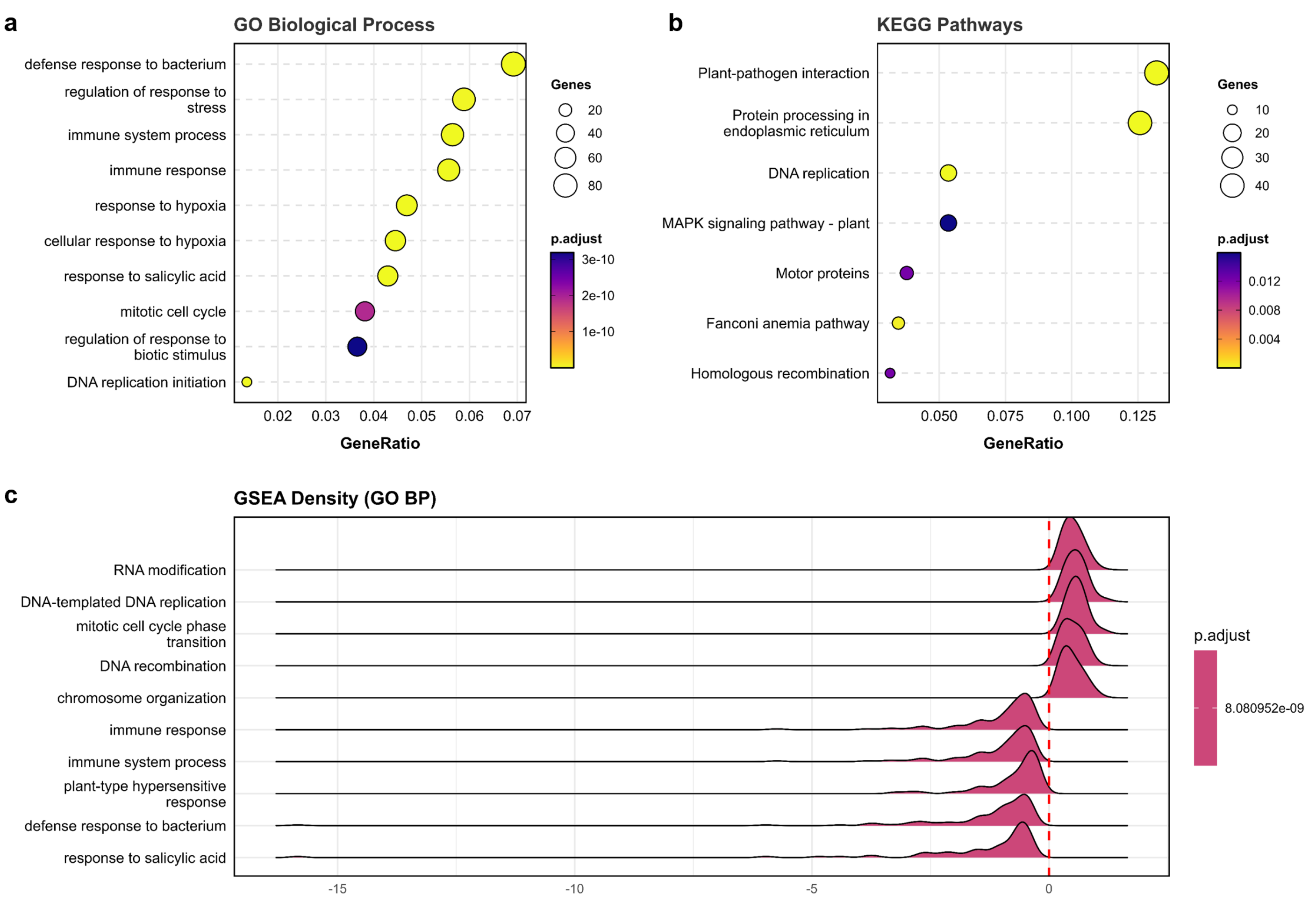


### Enrichment Analysis: dapat_Heat.Recovery_vs_Control.Recovery

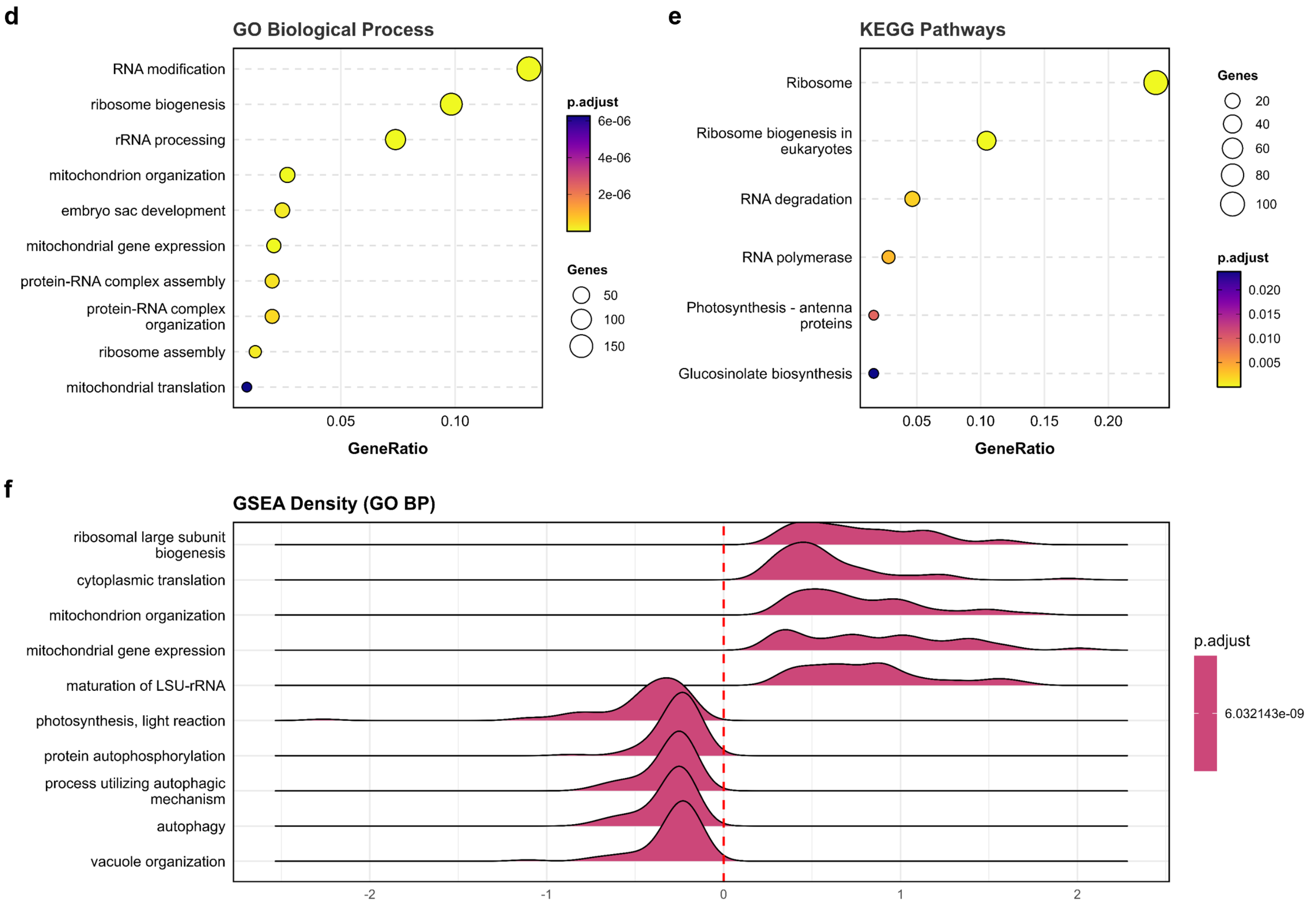
